# Herpes-like viral elements and universal subtelomeric ribosomal RNA genes in a chromosome-scale thraustochytrid genome assembly

**DOI:** 10.1101/2023.06.15.545109

**Authors:** Jackie L. Collier, Joshua S. Rest, Lucie Gallot-Lavallée, Erik Lavington, Alan Kuo, Jerry Jenkins, Chris Plott, Jasmyn Pangilinan, Chris Daum, Igor V. Grigoriev, Gina V. Filloramo, Anna M. G. Novák Vanclová, John M. Archibald

**Author notes:** these authors contributed equally (Jackie L. Collier, Joshua S. Rest).

## Abstract

We used long-read sequencing to produce a telomere-to-telomere genome assembly for the heterotrophic stramenopile protist *Aurantiochytrium limacinum* MYA-1381. Its ∼62 Mbp nuclear genome comprises 26 linear chromosomes with a novel configuration: subtelomeric rDNAs are interspersed with long repeated sequence elements denoted as LOng REpeated - TElomere And Rdna Spacers (LORE-TEARS). These repeats may play a role in chromosome end maintenance. A ∼300 Kbp circular herpesvirus-like genomic element is present at a high copy number. A 269 Kbp related virus-like element was found to reside between two complete sets of rRNA and LORE-TEAR sequences on one end of chromosome 15, indicating recent recombination between the viral and nuclear genome. Our data reveal new types of giant endogenous viral elements originating from herpes-like viruses and existing as either ‘stand-alone’ or integrated elements.

## MAIN

Labyrinthulomycetes are ecologically important osmoheterotrophic marine stramenopiles which possess interesting biochemical and cell biological novelties (Fossier Marchan et al. 2018). For example, the possession of a polyketide synthase-like fatty acid biosynthesis pathway has made some strains workhorses for production of the essential polyunsaturated fatty acid docosahexaenoic acid, and they may also be suitable for large-scale production of isoprenoid compounds like squalene and carotenoids (Rius et al. 2023; X. Xu et al. 2020). Labyrinthulomycetes were the source of pioneering discoveries in molecular biology, with notable examples published in *Science*. In the 1960s, Moens and Perkins (1969) demonstrated that *Labyrinthula* has a sexual cycle, haploid number of 9 chromosomes, and that the chromosome ends are associated with the nuclear membrane by tracing the synaptonemal complexes of prophase nuclei in serial sections by electron microscopy. In the 1970s, the first herpes-like virus detected outside of vertebrates was reported by Kazama and Schornstein (1972). Notably, neither of these important observations were followed up in the phylum in which they were discovered. The advent of long-read DNA sequencing (Rhoads and Au 2015; Branton et al. 2008) can rapidly advance the connection of old observations like these to the omic era by enabling the assembly of highly contiguous genomes. Chromosome-scale genomic sequences can reveal the abundance and distribution of repeat elements and ribosomal RNA (rRNA) gene clusters, the number and location of centromeres, the presence of endogenized viral genes and genomes, and the structure of telomeres and subtelomeric regions (Filloramo et al. 2021; Guérin et al. 2021; Fang et al. 2020; Fletcher et al. 2019; Moniruzzaman et al. 2020). Genome-scale leaps in biological insight can be particularly valuable for non-model organisms like the labyrinthulomycetes (Collier and Rest 2019), analysis of which can expand perspectives on eukaryotic biology. Here we present genome-wide discoveries in the labyrinthulomycete protist *Aurantiochytrium limacinum* ATCC MYA-1381 (a thraustochytrid) that corroborate and extend early reports about the genome structure and viral associates of labyrinthulomycetes.

## RESULTS AND DISCUSSION

### Chromosome Architecture and Subtelomere-Associated Elements in the *A. limacinum* Genome

We performed short-read 454 (454 Life Sciences) and long-read Nanopore (Oxford Nanopore Technologies (Jain et al. 2016)) sequencing of the *A. limacinum* nuclear genome, which independently yielded assemblies of 60.93 Mbp (Newbler) and 63.71 Mbp (Canu; (Koren et al. 2017)), respectively (Additional File 1: **Text S1, Fig. S1;** Additional File 2: **Table S1, Table S2, Additional File 3**). Neither assembly was rich in repetitive sequence, with ∼4% of the assemblies containing repetitive sequence (mostly simple repeats (Chen 2004) (Additional File 1: **Table S3**). Multiple metrics suggested that both assemblies were highly complete: 99.66% of RNA-seq reads mapped to the 454 assembly and 96% to the Nanopore assembly (Additional File 1: **Text S1**), and we detected 91.4% and 87.9% of Eukaryota BUSCO genes in the 454 and Nanopore assemblies, respectively (Simão et al. 2015) (8.6% and 12.1% missing BUSCOs, respectively; Additional File 1: **Table S4**).

We found that the 26 largest Nanopore contigs likely represent complete or nearly complete *A. limacinum* physical chromosomes. These contigs range in size from ∼1.02 Mbp to ∼4 Mbp (**Fig. 1**) and total 61.41 Mbp (96.4% of the complete Nanopore assembly); they align with 37 of the longest 454 scaffolds (referred to here as the primary 454 assembly) containing 59.93 Mbp (98.4%) of the total 454 assembly; Additional File 1: **Fig. S1B, Fig. S2;** Additional File 2: **Table S1, S2**). The Nanopore contig sizes are consistent with our examination of the genome by pulsed-field gel electrophoresis, which detected chromosome-sized bands ranging from ∼1.05 Mbp to >3 Mbp (Additional File 1: **Fig. S3**). This genome structure is similar to other stramenopiles for which both chromosome number and genome size are known: three diatoms and a eustigmatophyte have smaller genomes (31-36.5 Mbp) with similar numbers of chromosomes (24 to 33), while the oomycete *Phytophthora sojae* has an 82.6 Mbp genome with ∼12-14 chromosomes (Filloramo et al. 2021; Diner et al. 2017; Guérin et al. 2021).

**Fig. 1.**
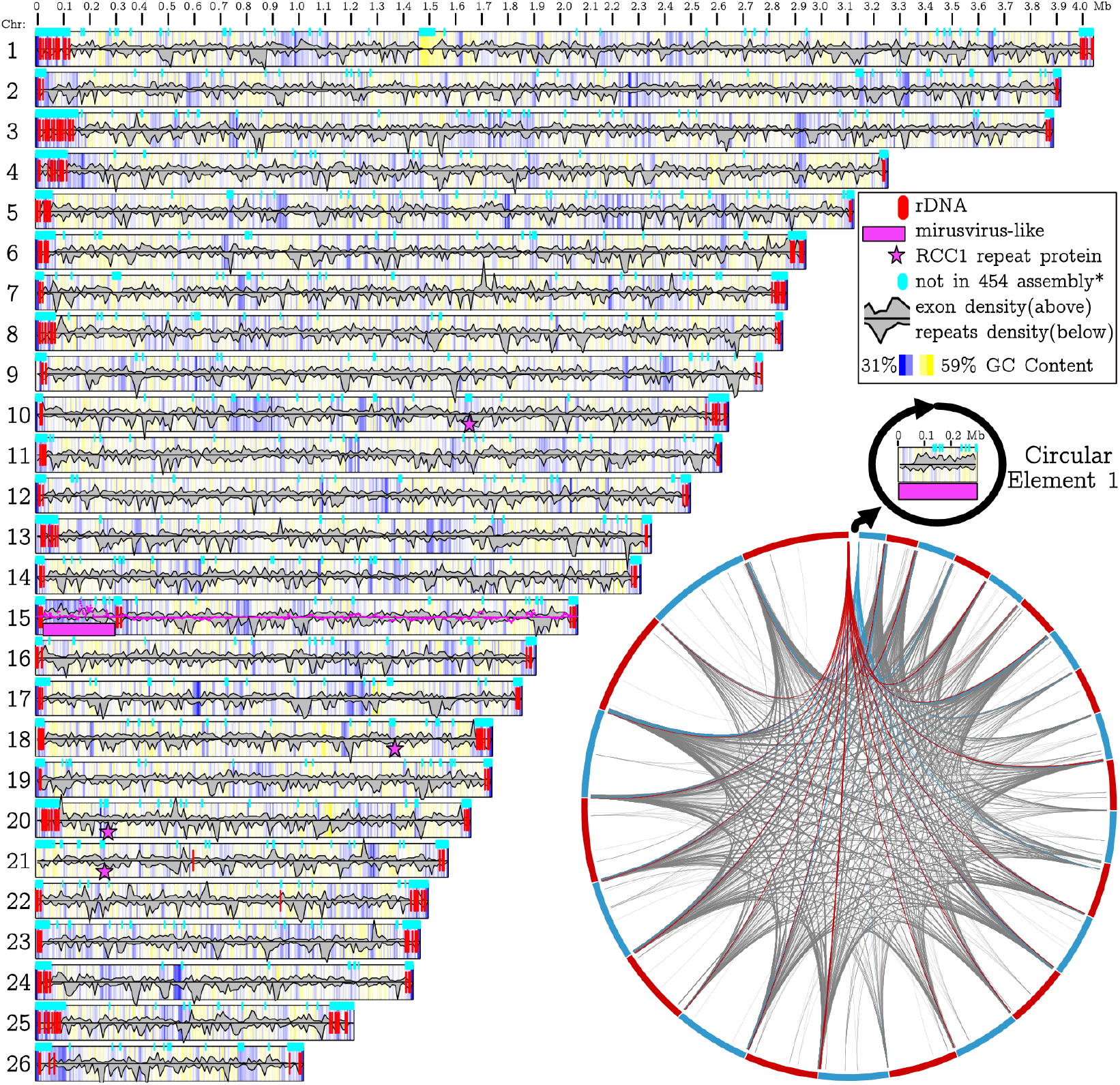
Size and select features of the 26 putative linear chromosomes and a Circular Element in *Aurantiochytrium limacinum* ATCC MYA-1381. Circular Element 1 is predicted to be circular, but is displayed as linear. A scale in megabases is provided along the top of the plot (Gel and Serra 2017). Vertical red lines represent locations of predicted (Lagesen et al. 2007) rRNA gene regions, and are found almost exclusively at the ends of the linear chromosomes. Cyan boxes represent regions that did not align (Darling et al. 2004) with the primary 454 assembly (*some regions may be present in short 454 scaffolds; see Supplemental Fig. S2). GC content (5 Kbp windows) is indicated by background color, with darker shades of blue indicating regions of lower GC content and darker shades of yellow indicating regions with higher GC content; low GC content at the linear chromosome ends reflect telomere content. The gray density plot above the midline of each chromosome indicates the relative density of exons, based on mapping of predicted exons from the JGI assembly to each Nanopore chromosome. The gray density plot below the midline (reflected so that higher values form valleys) indicates the relative density of repetitive sequences identified by RepeatMasker (Chen 2004). The magenta line (shown only for Chromosome 15) is a plot of the VirusRecall rolling score of NCLDV content (original range -17.6 to 3.9; negative and positive scores rescaled linearly above and below the axis, respectively). **Inset**: Chord plot showing matching sequence regions of at least 1 Kbp between contigs (Delehelle et al. 2018). An arbitrary set of chords are colored (arbitrarily red and blue) to highlight the directional nature of the repeats at the scaffold ends. A blank space in the chord plot represents the shortest scaffold, which has no matching sequence regions on other scaffolds.

Most *A. limacinum* chromosomes were sequenced telomere-to-telomere. Among the 52 predicted chromosome ends, 39 terminate with telomeric repeats of sequence TTAGG ∼500 bp in length (mean 480 bp, median 499 bp) (**Fig. 2**). The telomeric repeats identified in the Nanopore assembly are slightly shorter than the TTAGGG repeats of vertebrates and other eukaryotes, including some fungi, plants, and protists such as the photosynthetic stramenopile *Pelagomonas calceolata* (Guérin et al. 2021), but identical to TTAGG repeats reported from diverse insects and a few other eukaryotes (http://telomerase.asu.edu/sequences_telomere.html). Telomeric repeats are missing from 13 of the chromosome ends; this likely reflects assembly issues (Additional File 1: **Text S1**).

**Fig. 2.**
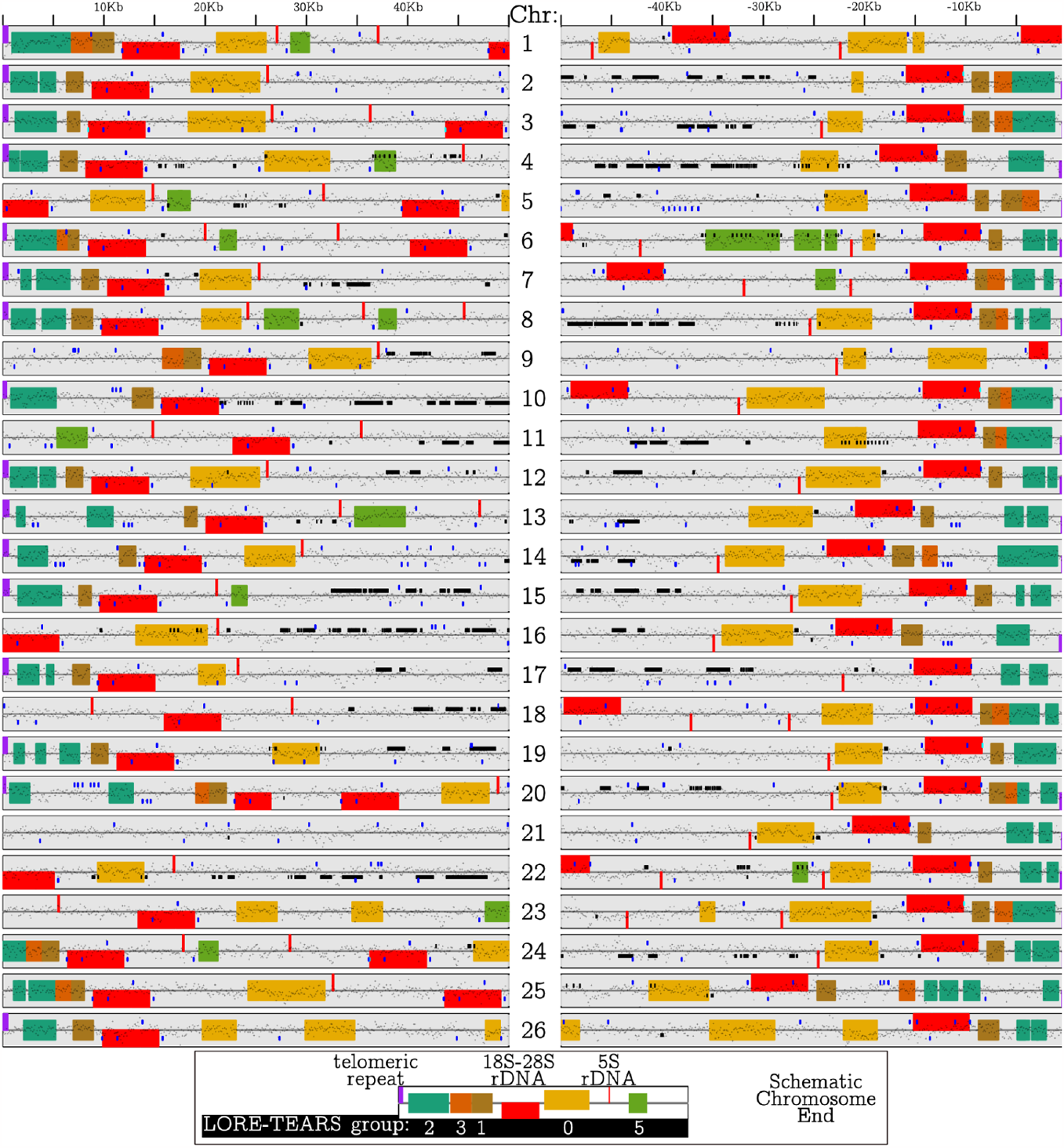
Focused view of the 50 Kbp of both ends of the *Aurantiochytrium limacinum* chromosomes (Chr) from Figure 1. Locations of predicted rRNA genes (red) as in Fig. 1, with a large 18S-5.8S-28S rDNA cluster transcribed toward the telomere, and a small 5S rDNA transcribed away from the telomere. Purple boxes on ends of contigs indicate the presence of telomeric repeats in our assembly. Five classes of LORE-TEARS (0,1,2,3,5) are shown (see inset for key). G-quadruplexes are plotted as blue lines; note the regular positioning of G-quadruplexes within and around the rRNAs. Black lines are exons mapped using BLAST. GC content is plotted along the centerline. **Bottom Inset:** Schematic view of a typical arrangement of elements at the ends of a chromosome. LORE-TEARS elements are colored and labeled by their sequence similarity group.

Almost all of the sub-telomeric regions of the *A. limacinum* chromosomes unexpectedly contain 18S, 5.8S, and 28S rRNA gene clusters interspersed with long repeated sequences (**Fig. 1, 2**). These elements are evident as extensive sequence matches between the ends of all contigs (**Fig. 1**, inset; in contrast to the 454 assembly; Additional File 1: **Fig. S4**). Specifically, in 37 of the 39 Nanopore contig ends with telomeres, an 18S-28S rRNA gene cluster (small subunit or 18S rRNA, ITS1, 5.8S rRNA, ITS2, large subunit or 28S rRNA; average length = 5551bp) transcribed toward the telomere is found ∼9.4 Kbp (median) from the telomeric repeat (**Fig. 2**). In 30 of these 37 contig ends, a 5S rRNA gene transcribed away from the telomeric repeat is found ∼10 Kbp (median) further from the telomere. In 17 of these 30 cases, no other rRNA genes were identified, and the 454 assembly scaffold mapped to the approximate location of the 5S gene. The remaining contig ends vary from this pattern. In some cases both ends of a contig have the same organization (Chr17, Chr15, Chr6), but more commonly the two ends are different. Only one rRNA gene (a 5S on Chr3) was found in the opposite orientation, and only three 5S genes (on Chr21, Chr22, Chr15) and one 18S-28S rRNA gene cluster (on Chr15) were identified in the more central regions of Nanopore assembly contigs (**Fig. 1)**.

Between each of these telomeric and subtelomeric elements are characteristic long repeated sequences (**Fig. 2;** Additional File 1: **Table S5;** identified with Tandem Repeats Finder (Benson 1999)). We call these LOng REpeated - TElomere And RDna Spacers (LORE-TEARS), which are built from 366 - 529 bp units repeated between 1.8 and 22.6 times and lacking similarity to sequences in GenBank. Several distinct LORE-TEARS families occur in regular positions with respect to the chromosome ends. Just downstream of the 28S rRNA genes, there is usually one ‘Group 1’ element containing ∼4 repeated units, each ∼406 bp long. Closer to the telomere, there is usually at least one ‘Group 2’ element with ∼6 repeated units, each ∼366 bp long. Usually upstream from the 18S gene nearest the telomere and between it and the nearest 5S gene is a ‘Group 0’ element, containing ∼9 repeated units of ∼385 bp. Where two consecutive 5S rRNA genes are detected, there is often a ‘Group 5’ element between them (∼5 repeats of a ∼421 bp unit). Among the 15 putative chromosomes with telomeric repeats assembled at both ends, seven have a ‘Group 3’ element (∼3 repeats of a ∼529 bp unit) between the ‘Group 1’ and ‘Group 2’ elements at only one end, while one has a ‘Group 3’ element between the ‘Group 1’ and ‘Group 2’ elements at both ends and seven have no ‘Group 3’ elements. We detected G-quadruplexes associated with rRNA genes and some LORE-TEARS, which is consistent with a regulatory function for these elements at the chromosome ends (Juranek and Paeschke 2012; Paeschke et al. 2008; Biffi, Tannahill, and Balasubramanian 2012; Wang et al. 2012) (**Fig. 2**, Additional File 2: **Table S6**).

The organization of rRNA genes in the subtelomeres of the *A. limacinum* chromosomes suggests a consistent and specific relationship with telomeric processes. This arrangement is highly unusual, both in the nature of the repeats and their location. Eukaryotic rRNA genes are most commonly organized in a few large tandem arrays (e.g., one in yeast, five in humans) often associated with sites of genomic fragility, and 5S genes and 18S-28S gene clusters are typically not associated with one another (Kobayashi 2011; Torres-Machorro et al. 2009). Subtelomeric rRNA tandem repeats have been found in plants (Dvořáčková, Fojtová, and Fajkus 2015; Roa and Guerra 2012) and in some metazoans (in aphids at one telomere of the X chromosome (Criniti et al. 2009)), and in the protist parasite *Giardia* (Tůmová et al. 2015; F. Xu, Jex, and Svärd 2020). The multicellular stramenopile *Saccharina japonica* (the kelp kombu) has a typical tandem 45S array in the middle of one chromosome, and a tandem 5S array at the subtelomere of another (L. Liu et al. 2017). In all these cases, rRNA gene arrays reside on the ends of only one or a few chromosomes. Unlinked 18S-28S rRNA gene clusters (i.e., not in tandem arrays) are found in the red alga *Cyanidioschyzon merolae* (Maruyama et al. 2004; Matsuzaki et al. 2004) and in several apicomplexan parasites (Torres-Machorro et al. 2010) including *Plasmodium falciparum* (Gardner et al. 2002). The 5S and 18S-28S coding regions in the Nanopore assembly of *A. limacinum* are more closely spaced than is usual for organisms where this linkage occurs, but not as tightly linked as in the brown alga (stramenopile) *Scytosiphon lomentaria* and some other protists, where the 5S is just downstream of the 18S-28S (Kawai et al. 1997, 1995).

The most similar sub-telomeric chromosome architecture to that of *A. limacinum* is found in the microsporidian parasites *Encephalitozoon cuniculi* and *E. intestinalis*, which have one subtelomeric, divergently transcribed 18S-28S rRNA gene cluster near the end of each of their 11 chromosomes separated from the telomeric repeats by two types of telomere-associated repeat elements (TAREs) with ∼30 to 70 bp repeat units (Mascarenhas Dos Santos et al. 2023). However, the 5S rRNA genes are not subtelomeric in *Encephalitozoon* spp. Subtelomeric 18S-28S rRNA gene clusters are also a chromosomal feature of the endosymbiotically-derived ‘nucleomorph’ genomes of cryptomonads (Douglas et al. 2001; Kim et al. 2022) and chlorarachniophytes (Suzuki et al. 2015). Brugère et al. (2000) suggested that the subtelomeric location of rDNA might be related to selective pressure associated with genome reduction, but the ∼62 Mbp genome of *A. limacinum* is not notably small among free-living stramenopiles, suggesting that genomic streamlining is not a factor here. The ends of chromosomes tend to be different from internal portions in exhibiting a higher frequency of recombination (Jensen-Seaman et al. 2004; McKim, Howell, and Rose 1988), lower level of gene expression (T. Liu et al. 2011), and higher rate of sequence evolution (Perry and Ashworth 1999). The selective forces and molecular mechanisms (e.g., convergent evolution by frequent inter- and intra-chromosomal homogenization) acting to maintain the consistent structure and homogeneous rDNA and LORE-TEARS sequences at the chromosome ends of *A. limacinum* offer novel avenues for future research, particularly if similar arrangements are found broadly in labyrinthulomycetes or other unexplored corners of protist diversity. The rRNA genes, LORE-TEARS, and/or associated subtelomeric sequences in *A. limacinum* may be involved in chromosome end maintenance and replication, including the maintenance of rDNA stability and/or nucleolar structure (Torres-Machorro et al. 2009), comparable to the repetitive subtelomeric sequences that are functionally important in other species (Tashiro et al. 2017; Scherf, Figueiredo, and Freitas-Junior 2001).

### Discovery of herpes-like viral elements in *A. limacinum*: Characterization of CE1 and LE-Chr15

We also detected a 298 Kbp chromosome of probable viral ancestry in *A. limacinum*, dubbed CE1 (circular element 1) (**Fig. 1**), with several remarkable features. This 27th genomic element is present in both genome assemblies (Additional File 2: **Table S1, S2**) and is consistent with a ∼0.35 Mbp band in the pulsed-field gel electrophoresis (Additional File 1: **Fig. S3)**. CE1 is predicted to be circular (Additional File 1: **Fig. S5**), and has read coverage (reads/bp) ∼9X higher than the other chromosomes, suggesting that it is present at a much higher copy number (Additional File 2: **Table S1**). CE1 lacks the predicted rRNA genes, LORE-TEARS, and telomeric repeats found on the other chromosomes (**Fig. 1**). GC content and mapped transcript abundance are similar to other chromosomes (Additional File 2: **Table S1**), but a smaller proportion of predicted genes have functional annotations, orthologous gene assignments, and predicted introns, and CE1 contains no BUSCO proteins (Additional File 1: **Fig. S6**). Of the 177 predicted genes on CE1, 128 are ORFans (i.e., do not hit any known proteins), 21 have best BLAST hits to bacteria, 22 to eukaryotes, four to archaea and one to viruses (when excluding the thraustochytrid *Hondaea fermentalgiana*; see below) (Additional File 2: **Table S7**).

VirSorter2 (Guo et al. 2021) and ViralRecall (Aylward and Moniruzzaman 2021) were initially used to identify both CE1 and the left end of Chr15 (LE-Chr15) as possible nucleocytoplasmic large DNA viruses (NCLDV) of the *Nucleocytoviricota* (Additional File 1: **Text S2, Table S8, Table S9**). Subsequent detailed sequence similarity searches using specific virion proteins of various groups of viruses as queries (via blastp and HMMsearch) against the *A. limacinum* genome revealed more genes on CE1 and LE-Chr15 related to key genes of herpes-like viruses recently identified as ‘*Mirusviricota*’ (Gaïa et al. 2023) than to core genes of *Nucleocytoviricota* (**Fig. 3A**; Additional File 2: **Table S7**; Additional File 1: **Text S2**). For example, we detected *Mirusviricota*-like major capsid protein (MCP) coding regions on CE1 and LE-Chr15 but no *Nucleocytoviricota*-like MCPs. A terminase-like homolog was also found to be shared between mirusviruses, CE1, and LE-Chr15 (the terminase protein packs the freshly synthesized genome into newly formed capsids). Divergent mirusvirus-like homologs were detected on both CE1 and LE-Chr15 for the remaining virion proteins as well (i.e., capsid maturation protease, portal protein, triplex 1 and triplex 2), as were other core mirusvirus genes, including a heliorhodopsin (on CE1), a histone H3 (CE1), TATA-binding protein (CE1 and LE-Chr15), subunits alpha and beta of ribonucleotide reductase (LE-Chr15), and additional proteins of unknown function (**Fig 3A;** Additional File 2: **Table S7;** Additional File 1: **Text S2**). We also detected core informational viral genes such as PolB (identified previously by Gallot-Lavallée and Blanc (2017), RNAPolB large subunit (RNAPL), and superfamily II helicase proteins (**Fig. 3A;** Additional File 1: **Table S7**). We found no sequence similarity between CE1 or LE-Chr15 and the lytic large DNA virus previously reported to infect the thraustochytrid *Sicyoidochytrium minutum* (SmDNAV) (Takao et al. 2007; Murakoshi et al. 2021) (Additional File 1: **Text S2**).

**Fig. 3:**
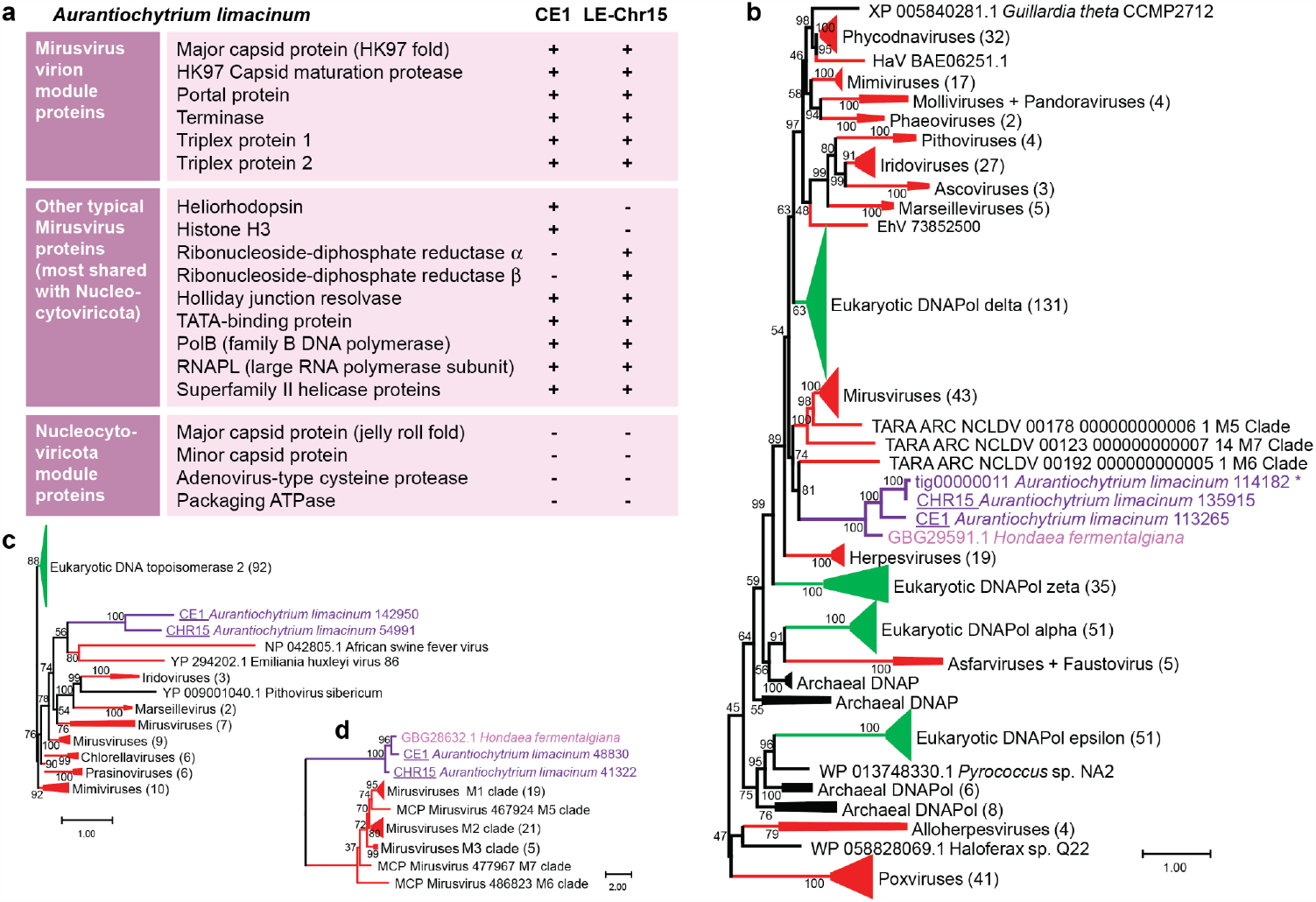
Viral content detected in *Aurantiochytrium limacinum* predicted proteins. **A**). Presence (+) and absence (-) of select viral proteins on CE1 and LE-Chr15 relative to key *Mirusviricota* and *Nucleocytoviricota* (NCLDV) proteins. Note that the RNAPL coding region is split into two discrete ORFs, as in pithoviruses and many archaea. **B**) Maximum likelihood phylogenetic tree of virus-like family B DNA polymerases (DNAPol) proteins encoded on CE1 and LE-Chr15 of *A. limacinum* (purple) and homologs in *Hondaea fermentalgiana* (pink). The sequences were aligned with MAFFT, and sites with less than 20% gaps were retained for phylogenetic reconstruction. Note that in addition to the homologs found in CE1 and LE-Chr15, a viral-like DNAPol is also found in *A. limacinum* tig00000011; it shows signs of pseudogenization. **C)** Phylogeny of viral DNA topoisomerases rooted with eukaryotic homologs. Sequences were aligned with MAFFT-linsi prior to phylogenetic reconstruction. **D)** Phylogenetic tree of Mirusvirus major capsid proteins and their homologs in *A. limacinum* and *H. fermentalgiana*. The sequences were aligned with MAFFT-linsi and sites with less than 30% gaps were retained for phylogenetic reconstruction. Viral sequences are in red, eukaryotic homologs are in green, and bacterial/archaeal sequences are in black. Scale bars indicate inferred number of amino acid substitutions per site.

Some virus-like genes on CE1 and LE-Chr15 are found only in mirusviruses or herpesviruses (MCP, **Fig. 3D**, terminases, **Fig. S7A**). For genes with broader distribution, phylogenetic analyses also support relationships of several CE1 and LE-Chr15 viral genes to mirusviruses, as well as to nucleocytoviruses, which share several informational genes with miruviruses. The resolvase, helicase, and DNAP trees show *A. limacinum* viral sequences branching specifically with mirusviruses (**Fig. 3B**; Additional File 1: **Fig. S7B and Fig. S7C**), while the topoisomerase and nuclease trees show relatedness of our *A. limacinum* viral sequences to nucleocytoviruses (**Fig. 3C; Fig. S7D**). In contrast, the *A. limacinum* viral TATA-binding proteins group with archaeal sequences, rather than mirusviruses (**Fig. S7E)**. Homologs of thraustochytrid viral and cellular arylsulfatase genes are detected only in various bacteria (**Fig. S7F**). The RNAPL genes of CE1 and LE-Chr15 are particularly unusual. As seen in some other viruses (e.g., *P. sibericum* (YP_009001268.1 and YP_009001052.1) and other pithoviruses, and cells (e.g., many archaea, (Langer et al. 1995)), the RNAPL coding region is split: the N- and C-terminal domains are encoded by separate ORFs located far apart from one another on both CE1 and LE-Chr15, and in *H. fermentalgiana*. The CE1, LE-Chr15, and *H. fermentalgiana* homologs branch robustly together in independent phylogenies of both RNAPL domains (**Fig. S7G and S7H**) but the precise evolutionary origin(s) of RNAPL in *A. limacinum* is unclear from the data in hand. On balance, these data suggest that CE1 and the viral-like element of Chr15 are most closely related to mirusviruses described in (Gaïa et al. 2023), but with genes derived from other sources as well.

The putative viral region at the left end of Chr15 (LE-Chr15) provides a nexus between the striking subtelomere structure and viral content of the *A. limacinum* genome. LE-Chr15 has rRNAs, LORE-TEARS, and telomeric repeats on one end and rRNAs and LORE-TEARs on the other (**Fig. 1, Fig. 2**); this is the only place in the assembly with internal (non-telomeric) arrays of rRNA genes and LORE-TEARs. The GC content of the putative viral integrant is 41.6%, slightly lower than the rest of the chromosome (45.0%), consistent with a foreign origin. The putative virus-like elements detected on CE1 and LE-Chr15 are related but distinct from one another (Additional File 1: **Fig. S8**). Comparing CE1’s 177 predicted proteins to the 152 proteins encoded by the virus-like region of Chr15, only 48 are each other’s reciprocal best BLAST hits (Additional File 2: **Table S7**), but many of their shared homologs branch together in phylogenetic trees (**Fig. 3B and D**; Additional File 1: **Fig. S7; Text S2**).

Mirusvirus virion particles have yet to be isolated. Our data show that *A. limacinum* is a probable natural host. CE1 could be an active viral genome capable of yielding viral particles: CE1 is circular and has an apparently complete virion module and full-length DNA polymerase, and viral particles consistent with the presence of an endogenous herpes-like virus have previously been identified in thraustochytrids (Kazama and Schornstein 1972, 1973). It is noteworthy that CE1 encodes proteins with ParA (Aurli_135839) and Fic (Aurli_13050) domains, which have been associated with plasmid segregation (Łobocka and Gagala 2020): this may speak to how CE1 is maintained as an episomal element in *A. limacinum*. Endogenization of LE-Chr15 appears to have occurred via sub-telomeric recombination. The pace of discovery of endogenous viral elements is accelerating thanks to growth of genome-scale resources in diverse organisms (Feschotte and Gilbert 2012; Schulz, Abergel, and Woyke 2022); to our knowledge, the differences between CE1 and LE-Chr15 make this the first example of related ‘stand-alone’ and integrated virus-like elements recovered through routine eukaryotic genome sequencing. Interestingly, we found several close homologs of the *A. limacinum* virus-like sequences in the fragmented genome assembly of another thraustochytrid, *Hondaea fermentalgiania* (Dellero et al. 2018) (**Fig. 3**, Additional File 1: **Fig. S7**). This suggests that mirusvirus-like viruses have been associated with this protist lineage for some time.

## Conclusions

Long-read sequencing has revealed that the genome of *A. limacinum* is dynamic and structurally innovative, an epic advance in light of their role as subjects first detailing chromosome counts in protists (Moens and Perkins 1969). We observed arrays of sub-telomeric rRNA and position-specific classes of long repeated elements (LORE-TEARS) on all chromosome ends, suggesting their role in chromosome maintenance. We also caught the ‘superposition’ of two complete sets of these elements surrounding an endogenized viral genome-like element at the end of Chr15. Our comparative genomic investigation reveals that CE1 and LE-Chr15 are specifically related to the recently discovered mirusviruses that are predicted to be “…among the most abundant and active eukaryotic viruses characterized in the sunlit oceans” (Gaïa et al. 2023). Mirusviruses have not, however, been linked to specific microbial eukaryotic hosts: Labyrinthulomycetes such as *A. limacinum* appear to be one such host. The functionality of both this integrated mirusvirus-like entity and of the circular, high-copy mirusvirus-like genome we identified are unclear, but both appear to correspond to (or be derived from) novel giant endogenous viral genomes, and one appears to maintain itself independently as a plasmid-like entity. It is noteworthy that Kazama and Schornstein described their thraustochytrid culture as ‘virogenic’ because they were never able to isolate strains free of the virus - even when establishing cultures from single zoospores - and because viral particles were only produced under permissive growth conditions (Kazama and Schornstein 1972, 1973). The combination of the union of highly conserved cellular elements (rRNAs), novel classes of repetitive elements (LORE-TEARs), and viral integration events at chromosome ends suggests new opportunities for future investigation of mechanisms of chromosome maintenance and nucleolus formation. There is still much eukaryotic diversity to be surveyed with long-read technology, and we will soon learn whether these features are unique to *A. limacinum* or a general feature of labyrinthulomycetes or broadly distributed among eukaryotic diversity.

## ONLINE METHODS

### Strain cultivation and nucleic acid preparation

*Aurantiochytrium* (formerly *Schizochytrium*) *limacinum* Honda et Yokochi ATCC MYA-1381 (also designated NIBH SR21 or IFO 32693, GenBank Accession AB022107; (Honda et al. 1998; Yokoyama and Honda 2007) was isolated from seawater in a mangrove area of the Yap Islands, Micronesia. For sequencing at JGI, *Aurantiochytrium* ATCC MYA-1381 cultures were grown in 2 liters of ATCC 790 By+ medium (5 g glucose, 1 g yeast extract, 1 g peptone, 30 g Instant ocean per liter) distributed in four large tissue culture flasks (500 ml each) at room temperature without shaking. Cultures were harvested after 7 days, producing 2.4 g wet weight. Genomic DNA was extracted from 0.507 g wet biomass and RNA from 1.057 g wet biomass following the protocols of (Lippmeier et al. 2009) and subject to JGI QA/QC protocols. For Nanopore sequencing, *Aurantiochytrium* ATCC MYA-1381 was cultured for three days in 50 ml ATCC 790 By+ medium. Genomic DNA was extracted based on a previously published protocol (https://dx.doi.org/10.17504/protocols.io.n83dhyn). The precipitated DNA was left to dissolve in water by spontaneous diffusion for 48+ hours at room temperature to avoid shearing and subsequently purified using QIAGEN Genomic-tip 20/G. Agarose gel electrophoresis (1%) was used to visually assess and confirm the integrity of high molecular weight (20+ Kb) DNA. DNA quality was evaluated using a NanoPhotometer P360 (Implen) to measure A260/280 (∼1.8) and A260/230 (2.0-2.2) ratios. The quantity of DNA was calculated using a Qubit 2.0 Fluorometer (ThermoFisher Scientific) with the dsDNA broad range assay kit.

### Sequencing and assembly

Short-read sequencing was performed by the Joint Genome Institute on the 454 sequencing platform, and assembly was accomplished with Newbler followed by annotation with the JGI Annotation Pipeline; details are provided in Additional File 1: **Text S1**. Draft genome sequence for *Aurantiochytrium limacinum* ATCC MYA-1381 is available via the PhycoCosm Genome Portal, (https://mycocosm.jgi.doe.gov/Aurli1/Aurli1.info.html).

Long-read sequencing was performed using the Oxford Nanopore Technology (ONT) MinION sequencing platform and assembly was accomplished with Canu; details are provided in Additional File 1: **Text S1**. The raw fast5 MinION data has been deposited in the NCBI SRA database BioProject PRJNA680238 (WT accession: SRR13108467; KO32 accession: SRR13108466; KO33 accession: SRR13108465).

### rRNA, G-quadruplex, and repetitive element predictions

rRNA gene locations were predicted using RNAmmer 1.2 (Lagesen et al. 2007). Tandem Repeat Finder 4.09 (Benson 1999) was used to identify tandem repeats (TR) with maximum repeat unit set as large as possible (2000 bp). The vast majority of the 29640 identified TRs were a few bp in length, and the number of TRs declined with TR length until ∼350 bp, where a peak appeared. The sequences of the 673 TR elements longer than 299 bp were dereplicated by removing overlapping TRs, keeping the shorter repeat unit and clustering with cd-hit. Telomeric repeats in the Nanopore assembly were identified in this output as repeats with unit 5, 10, or 15 matching the motif TTAGG (or CCTAA). More than 98% of TR elements were shorter than 200bp. 490 Tandem repeat (TR) units longer than 200bp were dereplicated (retaining one TR to represent each locus) and 398 subjected to clustering by cd-hit-est (Huang et al. 2010) with default parameters except sequence identity cutoff 0.8 and -r yes. Manual examination and alignment of the resulting 25 clusters (which excluded 34 singletons) revealed 4 types of TRs grouped by location relative to rRNA genes and telomeric repeats. Additional repetitive content was identified with RepeatMasker (Chen 2004). G-quadruplexes were predicted with G4-iM Grinder (Belmonte-Reche and Morales 2020) using Methods 2 and 3 and filtering for scores greater than 20. Genomic features were visualized using karyoploteR (Gel and Serra 2017).

### Viral gene predictions and phylogenetics

Virion proteins from various groups of dsDNA viruses, including mirusviruses and nucleocytoviruses, were used to screen the *A. limacinum* assembly using blastp and HMMsearches. HMMs were generated from alignments in Gaïa *et al*. (2023) (Additional File 3). Following the detection of mirusvirus-like structural proteins in CE1 and LE-Chr15, all mirusvirus ORFs (>=90 amino acids; predicted from the mirusvirus contigs from (Gaïa et al. 2023)) were used as blastp queries to detect additional mirusvirus homologs. In parallel, ViralRecall (Aylward and Moniruzzaman 2021) and VirSorter2 (Guo et al. 2021) were used to evaluate viral gene content in both the 454 and Nanopore *A. limacinum* assemblies. The results of similarity searches against nr were then used to characterize additional CE1 and LE-Chr15 proteins.

Unless specified otherwise, *A. limacinum* virus-like proteins were aligned with homologs in diverse viruses, prokaryotes and eukaryotes using MAFFT, with BMGE (default parameters) used to perform site selection. Maximum likelihood phylogenetic trees were constructed with IQTree (Nguyen et al. 2015) v1.6.3 model C60+G4 with 1000 ultrafast bootstraps replicates.

### Availability of data and materials

The short-read (454) data, assembly, and annotation are publicly available via the JGI genome portal in PhycoCosm (https://phycocosm.jgi.doe.gov/). The raw fast5 MinION data has been deposited in the NCBI SRA database BioProject PRJNA680238 (WT accession: SRR13108467; KO32 accession: SRR13108466; KO33 accession: SRR13108465).

Additional File 1 contains supplemental text, tables, and figures.

Additional File 2 contains supplemental tables S1, S2, S6 and S7.

Additional File 3: The Canu nanopore assembly, annotations, HMM files, alignments, and trees, are at doi:10.5061/dryad.2fqz612t6: https://datadryad.org/stash/share/QYfnoRY6ZF9b-xA1lCXvm7xfAjmeqY-A5b-nSv15YNI

## Authors’ contributions

JC and JR conceived the study, performed the analyses and drafted the manuscript. EL built the combined Canu assembly and performed initial comparative analyses. CD, JJ, JP, CP, AK, IVG performed the short-read sequencing, assembly, and annotation. GF and AMGNV carried out Nanopore long-read sequencing and performed the PFGE. JC, JR and LG-L performed detailed comparative genomic analyses of virus-like sequences, and LG-L and JMA performed and interpreted phylogenetic reconstructions. All authors read and approved the final manuscript.

## Acknowledgements

Thanks to Bruce A. Curtis for technical support with Nanopore sequencing and feedback on the final manuscript.

