## Additional File 1 for "Herpes-like viral elements and universal subtelomeric ribosomal RNA genes in a chromosome-scale thraustochytrid genome assembly"


Text S1: Details of Genome Assemblies
- Short-read genome and transcriptome sequencing, assembly, and annotation
- Long-read genome sequencing, assembly, and annotation
- Comparison of short-read and long-read assemblies

Text S2: Viral Content in the Aurantiochytrium limacinum genome
- CE1 has mirusvirus ancestry
- Chromosome 15 has sub-telomeric viral content bounded by rRNAs
- Viral elements in other labyrinthulomycetes and stramenopiles
- CE1 is smaller than expected.
- Potential virophage and other viral DNA in A. limacinum genome

Table S1. Nanopore assembly information.
Table S2. 454 Assembly information.
Table S3: RepeatMasker Output
- 454 Assembly
- Nanopore Assembly

Table S4. BUSCO 5.3 results from dataset eukaryota_odb10 on the Nanopore and 454 genome assemblies.
Table S5.  LOng REpeated TElomere And Rrna Spacers (LORE-TEARS) statistics
Table S6. G-quadruplexes detected within 28S rDNA genes
Table S7: Viral content detected in A. limacinum predicted proteins by HMM and BLAST.
Table S8: Viral (NCLDV) sequences identified by ViralRecall
Table S9: Viral sequences identified by VirSorter
Figure S1. Mauve mapping
Figure S2. Mapping between Nanopore and 454 Assemblies
Figure S3. Physical chromosome sizing
Figure S4. ASGART comparison
Figure S5. Nanopore tig00000136 assembly and 454 scaffold_34 assembly dotplots
Figure S6. Proportion of genes missing annotations across scaffolds
Figure S7. Phylogenetic trees of select proteins encoded on CE1 and the LE-Chr15 viral-like element

Fig. S7A. Phylogeny of terminases rooted with herpesviruses homologs.
Fig. S7B. Phylogeny of Holliday junction resolvases.
Fig. S7C. Phylogeny of superfamily 2 helicases.
Fig. S7D. Phylogeny of PD-DEXK nucleases.
Fig. S7E. Phylogeny of TATA box-binding proteins.
Fig. S7F. Phylogeny of arylsufatases.
Fig. S7G. Phylogeny of RNAPol N-terminal subunit.
Fig. S7H. Phylogeny of RNAPol C-terminal subunit.
Figure S8. tBLASTx comparison of viral regions on CE1 and chromosome 15 (LE-Chr15)
References

### Text S1: Details of Genome Assemblies

#### Short-read genome and transcriptome sequencing, assembly, and annotation

Short-read sequencing, assembly, and annotation of the *Aurantiochytrium* (formerly *Schizochytrium*) *limacinum* Honda et Yokochi ATCC MYA-1381 genome was performed at JGI. The genome was then annotated with the aid of the assembled transcriptome.

The genome was initially assembled from 454 (454 Life Sciences) reads generated from both standard unpaired and paired-end libraries. For the unpaired 454 Titanium Rapid library, genomic DNA samples were fragmented via sonication to 400-800 base pairs (bp). These fragments were end-polished and ligated to a set of 454 Y-shape adaptors. The 454 library fragments were then clonally amplified in bulk by capturing them through hybridization on microparticle beads and subjecting them to emulsion based PCR, resulting in beads that were covered with millions of copies of a single DNA fragment (range 400-800 bp) and where each bead contained a different clonally amplified library fragment. After amplification, the beads were recovered from the emulsions and loaded into the wells of a PicoTiterPlate device (PTP) such that wells contained single DNA beads. The PTP was then inserted into the 454 Genome Sequencer FLX-Titanium instrument for sequencing, where sequencing reagents were sequentially flowed over the plate and the sequence of the DNA fragments was determined.

For the 454 Titanium Paired-end (PE) library, 15 µg of genomic DNA was sheared by a Hydroshear to ~8 kilobase pair (Kbp) size fragments. The sheared sample was then gel selected for intense 8 Kbp bands, purified, and ligated to 42 bp loxP linkers on either end. These loxP linkers were labeled with biotin, then circularized by the Cre recombinase. As a result, the ends of 8 Kbp fragments were brought together and bridged by a single loxP linker. These circular DNAs were further sheared to 500 bp fragments and the fragments carrying the loxP linkers were recovered by Streptavidin-coated magnetic beads. 454 Titanium adaptors A and B were then ligated to the enriched loxP linker-containing fragments in the same way the unpaired libraries were created. The 454 library fragments were then clonally amplified and sequenced as with the unpaired library.

The short-read assembly was generated using Newbler version 2.6 (build:20110517_1502; (Silva et al. 2013). The accuracy of the assembly was assessed using 13 Sanger-sequenced fosmid clones. Fosmid DNA was isolated from a single bacterial colony and purified on a Qiagen MaxiPrep column. DNA was sheared to 3-4 Kbp using Adaptive Focused Acoustics technology (Covaris, Woburn, MA, USA) and cloned into the plasmid vector pIK96 as previously described (Ferris et al. 2010). Universal primers and BigDye Terminator

Chemistry (Applied Biosystems) were used for Sanger sequencing randomly selected plasmid subclones to a depth of 10x. The Phred/Phrap/Consed suite of programs were then used for assembling and editing the sequence (Ewing and Green 1998; Ewing et al. 1998; Gordon, Abajian, and Green 1998). After manual inspection of the assembled sequences, finishing was performed both by resequencing plasmid subclones and by walking on plasmid subclones or the fosmid clone using custom primers. All finishing reactions were performed using dGTP BigDye Terminator Chemistry (Applied Biosystems). Finished clones contained no gaps and were estimated to contain less than one error per 10,000 bp. All 13 fosmid clones were aligned to the assembly. In ten of the clones, the alignments were of high quality, and the overall bp error rate in this group of clones is 0.040% (152 bp discrepant out of 392,816 bp). Three clones (14866, 14868, and 14872) exhibited noteworthy discrepancies: clone 14866 stems from a repetitive region that spans a gap in scaffold00020 and has been collapsed in the assembly; the alignment of clone 14868 indicates that Newbler has inserted scaffolded gaps in the clone region; and clone 14872 falls into a repetitive region that appears to be collapsed in the assembly.

The transcriptome was used to assess the completeness of the genome assembly and to seed and assess the genome annotation. The transcriptome was sequenced using 454. A cDNA library was generated using the cDNA 454 Rapid Library Preparation Kit (Roche). mRNA was purified from total RNA using the Absolutely mRNA purification kit (Stratagene) and chemically fragmented using high heat. The fragmented RNA was reverse transcribed using random hexamers and AMV RT followed by second strand synthesis. The cDNA fragments were treated with end repair and ligated with 454 adapters. The 454 library fragments were then clonally amplified and sequenced as for genome sequencing. The resulting reads were assembled into RNA contigs using Newbler. To assess completeness, RNA sequences from one library (CHBC) were mapped to the genome assembly. Of 1509263 total EST sequences, only 60303 (4.0%) were not found, suggesting that the assembled genome is not missing large transcribed portions.

The 454-based JGI assembly contained 1662 contigs (N50/L50 233/82.5 megabase pairs (Mbp)) in 181 scaffolds (N50/L50 10/2.5 Mbp) with 937 assembly gaps (1.5% of the total 60.93 Mbp scaffold length). Scaffold lengths are shown in Additional File 2: **Table S2**. The genome was annotated using the JGI Annotation Pipeline, which detects and masks repeats and transposable elements, predicts genes, characterizes each conceptually translated protein with sub-elements such as domains and signal peptides, chooses a best gene model at each locus to provide a filtered working set, clusters the filtered sets into draft gene families, ascribes functional descriptions (such as GO terms and EC numbers), and creates a JGI genome portal in PhycoCosm (https://phycocosm.jgi.doe.gov/) with tools for public access and community-driven curation of the annotation (Kuo, Bushnell, and Grigoriev 2014; Grigoriev et al. 2021).

#### Long-read genome sequencing, assembly, and annotation

A multiplexed Nanopore (MinION, Oxford Nanopore Technologies) sequencing library for the wild-type *Aurantiochytrium limacinum* Honda et Yokochi ATCC MYA-1381 (WT) genome and two putative *crtIBY* knockout mutants (designated KO32 and KO33; (Rius et al. 2023)) was prepared using the Oxford Nanopore Technology (ONT) ligation sequencing kit (SQK-LSK109) and the PCR-free native barcoding expansion kit 1-12 (EXP-NBD103) according to the Oxford Nanopore Technologies protocol "1D Native barcoding genomic DNA with EXP-NBD103 and SQK-LSK109" (version NBE_9065_v109_revB_23May2018). Approximately 2 µg of purified genomic DNA per sample were used as input. Unfragmented genomic DNA for the wild-type and putative knockouts was repaired using the NEBNext FFPE DNA repair module (NEB cat. no. M6630) and prepared for adapter ligation using the NEBNext End repair/dA-tailing module (NEB cat. no. E7546) with incubations at 20°C and 65°C for 10 minutes each. The DNA repaired/end-prepped samples were purified with a 1:1 volume of AMPure XP beads (Beckman), and subjected to an incubation at room temperature for 10 minutes; the pelleted beads were subsequently washed twice with 80% ethanol. The DNA was eluted off the beads in 25 µl nuclease free water for 10 minutes at 37°C to encourage the elution of long molecules from the beads. The native barcodes NB07, NB08, and NB09 were ligated to the WT, KO32, and KO33 repaired/end-prepped DNA samples, respectively, in a 1.36x scaled ligation reaction. Each native barcoded sample was pooled in approximately equimolar amounts (~1.3 µg each). The 1D barcode sequencing adapters (BAM 1D) were then ligated to the pooled and barcoded DNA using a 1-hour incubation at 25°C. The adapter ligated DNA was purified by a 0.4x AMPure XP bead clean-up including a 10 minute incubation at room temperature and two washes using the Long Fragment Buffer mix to enrich for DNA fragments >3 Kbp. The final adapter ligated library was incubated in 15 µl Elution Buffer for 10 minutes at 37°C. A total of 1.2 µg of prepared library was loaded on a single MinION R9.4.1 chemistry SpotON flow cell (FLO-MIN106) and sequenced via Oxford Nanopore Technology's MinKNOW software (v2.1.12) without live basecalling. The raw fast5 MinION data been deposited in the NCBI SRA database BioProject PRJNA680238 (WT accession: SRR13108467; KO32 accession: SRR13108466; KO33 accession: SRR13108465).

Long-read genome assembly was performed on each of the three strains individually (described in (Rius et al. 2023)), and in combination.

**Strain-specific assemblies** (see (Rius et al. 2023) for detailed methods)**:** Briefly, for wild-type, the genome assembly totaled 61.9 Mbp in 55 contigs, while the genomes of KO mutants 32 and 33 both assembled as 62.5 Mbp into 50 and 47 contigs, respectively. Analysis by Mauve (Darling et al. 2004) revealed ~27 homologous contigs among the three assemblies, ranging from ~0.35 up to ~4 Mbp, and the detection of putative telomeric repeats at one or both ends of many of these contigs suggested they represent nearly chromosome-scale assemblies. The locus at which the three strains differ (*crtIBY*; (Rius et al. 2023)) was identified on putative chromosome 23. The wild-type assembly was the least contiguous (8 broken putative chromosomes) and had one apparently chimeric chromosome compared to KO32 and KO33 (**Fig. S1A**), suggesting that its lower mean read length (4913 bp vs 8508 and 7951) resulted in reduced contiguity. All three assemblies contained a relatively high-copy, ~350 Kbp contig suggested by Canu to be circular, representing CE1.

**Combined assembly:** In an effort to resolve the differences among these assemblies and take advantage of greater coverage, reads for all three strains were concatenated into one file as input for Canu v1.7.1 (https://github.com/marbl/canu), and read trimming and assembly were performed with default settings and an estimated genome size of 60 Mb. This combined Nanopore-based Canu assembly contained 62 contigs and a total of 63.71Mbp (**Fig. S1A**).The 27 contigs described in the main text total 61.77 Mbp. The remaining 35 contigs included five putative mitochondrial contigs and another 30 contigs ranging from 20 Kb to 230 Kb, of which 14 represented single reads (Additional File 2: **Table S1**). To assess completeness, the same RNAs used to assess the completeness of the 454 assembly were also mapped to this Nanopore assembly. Of 1509263 total RNA sequences, only 4395 (0.3%) were not found, suggesting that the Nanopore assembly is even more complete than the 454 one. This combined assembly was the focal point for the genome-scale investigations described herein because it generally had the longest version of each putative chromosome and had the most contigs with telomeres at both ends.

**Distribution of chromosome lengths:** In diverse eukaryotes, a typical distribution of chromosome lengths has been described (Li et al. 2011). Overall the observed distribution of the 17 longest *A. limacinum* Nanopore contig lengths is consistent with the expected distribution described in Li *et al.* (2011), with the exception that the 27th element (CE1) is smaller than expected relative to the other chromosomes we identified in *A. limacinum* (see below image.)

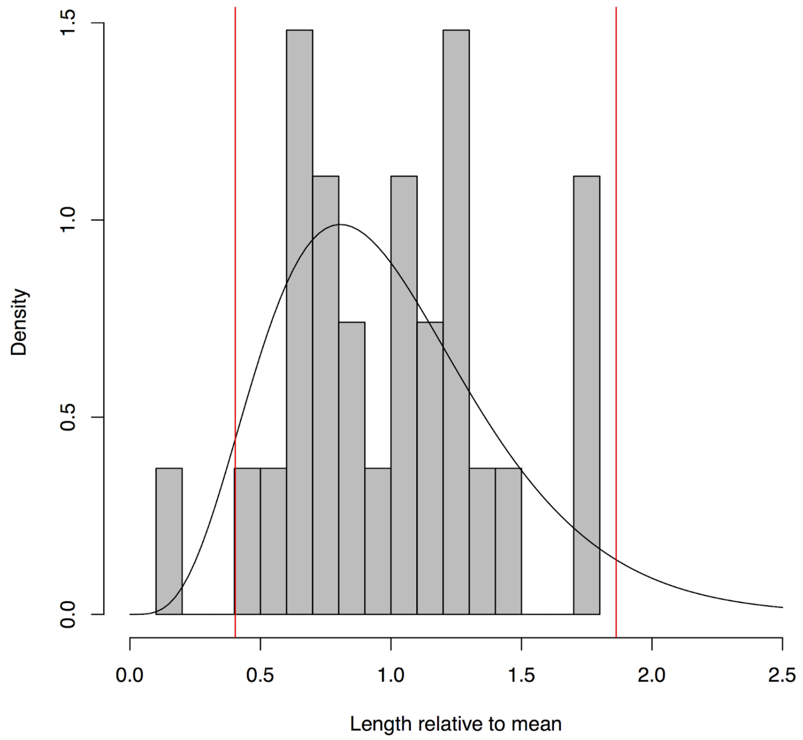


***Distribution of chromosome lengths*** *(Canu), expressed as a ratio to the mean length. A gamma distribution fit to the data is plotted as a black line; chromosome lengths typically follow this distribution (Li et al. 2011). The expected minimum (0.4035) and maximum (1.8626) ratios are plotted in red, as predicted by Li et al. (2011). Tig00000136 (CE1) is the histogram bar farthest to left, with a ratio of 0.16, and falls outside of this expectation.*

#### Comparison of short-read and long-read assemblies

We used Mauve to align the 454 and Nanopore assemblies. Mauve aligned the 27 longest Nanopore contigs with the 38 longest 454 scaffolds, which ranged in size from ~0.148 Mb to ~5.48 Mb and contained 60.22 Mb (98.8% of the total 454 assembly). In 16 cases, elements from these two independent assemblies aligned one to one (Additional File 2: **Table S2; Fig. S1B**), with the 454 scaffold slightly shorter (2% on average) than the matching Nanopore contig. Two additional Nanopore contigs (4065 and 4069) were each >98% covered by a pair of 454 scaffolds. The alignments between the remaining 10 contigs and 19 scaffolds were less straightforward, as five 454-based scaffolds were split between two or more Nanopore contigs (Additional File 2: **Table S2)**. The 454 assembly's scaffold_1, which was the longest in that assembly at 5.475 Mb in length and longer than any Nanopore contig, mapped to three Nanopore contigs: ~3.1 Mb with tig0000318, ~2.3 Mb to tig0000010, and a small part to tig0000088. It seems likely that a ~20 Kb assembly gap combined parts of two separate chromosomes represented by tig0000318 and tig0000010 into scaffold 1. Similarly, 454 scaffolds 7 and 23 were split fairly evenly between two Nanopore contigs, and scaffold 18 split fairly evenly among three Nanopore Canu contigs. 454 scaffold 26 mapped mainly to tig00000106, with a small part to tig00000329. Overall, this comparison suggests a tendency for Newbler to assemble chimeric scaffolds.

The 30 shortest (<230 Kb) non-mitochondrial contigs in the Nanopore assembly contained the majority (82%) of the 4.84 Mb of the Nanopore-based assembly found in islands (>=20 bp) in the Mauve alignment of the 454 and Nanopore assemblies. A BLAST search of these contigs using the JGI scaffolds as a query indicates a continuous distribution of many short matches, and 29 of the 30 contain rRNA genes and telomeric repeats (Additional File 2: **Table S1**); we interpret these as representing mis-assembled / unassembled chromosome ends. The complementary comparison (454 islands not in the Nanopore assembly) was 2.6% (330 kbp), and many (24%) of the islands in the 454 assembly were from the ends of the 454 scaffolds, comprising poor quality sequence (homopolymer stretches, stretches of N's, etc.) that made it past the assembler threshold, but likely representing artifacts, consistent with previous analyses of 454 data (Huse et al. 2007; Archer et al. 2012). The 143 454 scaffolds not included in the main genome alignment are small, although all have one or, typically, many BLAST hits to Nanopore contigs, consistent with containing repetitive or difficult to align sequences (**Fig. S2**). These scaffolds contain a small number of predicted protein-coding genes (320 genes across the 142 short scaffolds).

We used ASGART (Delehelle et al. 2018) to visualize segmental duplications in the two assemblies. The Nanopore assembly contained many more matches than the 454 assembly (**Fig. S4**). The sequences ASGART detected as shared among Nanopore contigs corresponded to the presence of directional arrays of rRNA genes (predicted with rnammer (Lagesen et al. 2007)) near both termini of most Nanopore contigs (main text; **Fig. 1**, **Fig. 2**) and not present in the 454 assembly. The LORE-TEARS sequences that neighbor the rDNA are present in the secondary (i.e. short scaffolds) of the 454 assembly (main text and **Fig. S2**); the rDNAs themselves were masked from the 454 assembly (**Fig. S2B**).

### Text S2: Viral Content in the *Aurantiochytrium limacinum* genome

Notes: scaffold_35 (454) aligns with the left end of tig00004069 (nanopore), which we call chromosome 15 in the manuscript. rscaffold_34 aligns with tig00000136 (nanopore), which we call Circular Element 1 (CE1) in the manuscript.

#### CE1 has mirusvirus ancestry

CE1 was initially identified as an NCLDV by ViralRecall (Aylward and Moniruzzaman 2021) at the contig level (score=0.185 for scaffold_34 from the 454 assembly, and score=0.113 for tig00000136 from the Nanopore assembly) (**Table S8**). ViralRecall detected 0.84-0.88 viral hits for every non-viral (pfam) hit. ViralRecall also identified specific regions in CE1 that have NCLDV content; from the 454 assembly the region was 288 Kbp in length (out of 297 Kbp, score=0.205), and from the Nanopore the region was 247 Kbp in length (out of 298 Kbp, score=0.183). In these regions, between 10-14% of ORFs were categorized as viral. ViralRecall identified the viral marker genes PolB (family B DNA Polymerase) (Iyer, Aravind, and Koonin 2001) and RNAPL (the large RNA polymerase subunit) (Aylward et al. 2021; Moniruzzaman, Martinez-Gutierrez, et al. 2020) on CE1.

CE1 was also identified as an NCLDV by VirSorter2 (Guo et al. 2021) when scaffold_34 from the 454 assembly was analyzed (**Table S9**). VirSorter2 characterized the putative NCLDV region as 191 Kbp long (of 297 Kbp) with one hallmark gene (a putative superfamily II helicase); among all identified genes, 6.2% were categorized as viral and 1.2% were categorized as cellular. However, VirSorter2 did not identify tig00000136 from the Nanopore assembly as an NCLDV.

With the recent discovery of the mirusviruses (Gaïa et al. 2023), we revisited our ViralRecall and VirSorter2 results and performed a more focused analysis of the gene content of CE1 and LE-Chr15 relative to the known diversity viruses, prokaryotes and eukaryotes (note that mirusviruses are not presently included in databases used by ViralRecall and VirSorter2). Mirusviruses appear to share several of their informational genes with the Nucleocytoviricota (*Varidnaviria*), while their virion module is most closely related to Caudoviricetes (*Duplodnaviria*) (Gaïa et al. 2023). This helps explain the apparent evolutionary affinities of the virus-like elements in *A. limacinum* and explains why the viral sequences in *A. limacinum* originally seemed to be of NCLDV ancestry, given that the mirusviruses are not presently included in the software databases.

#### Chromosome 15 has sub-telomeric viral content bounded by rRNAs

NCLDV content was initially identified on Chr15 by ViralRecall; from the Nanopore assembly the region was 166 Kbp in length, spanning from positions 111653 to 278329 (score=0.442) (**Table S8**). This region contained 33 viral hits out of 239 ORFs. It contained two marker genes: RNAPL (the large RNA polymerase subunit) and ribonucleotide reductase (RNR). A short region (length 22 Kbp) was also identified by VirSorter2, although it was classified as a dsDNA phage, rather than an NCLDV, with a score of 0.887, 1 hallmark gene, 3.6% viral genes, and no cellular genes (**Table S9**). This region of chromosome 15 (see **Fig. S1**) corresponds to almost the entirety of scaffold_35 from the 454 assembly (except ~12 kbp at one end and ~9 kbp at the other, corresponding to regions with rRNA sequences in the matching Nanopore contig). ViralRecall categorizes 249 Kbp (out of 295 Kbp) of scaffold_35 as an NCLDV (score=0.620), with 16% viral gene content and  and three NCLDV marker genes: RNAPL, RNR, and PolB. Similarly, VirSorter2 categorizes a 273 Kbp region (out of 295

Kbp) of scaffold_35 as an NCLDV (score=0.727). (ViralRecall also categorizes the entirety of scaffold_35 as an NCLDV; score=0.512).

#### Viral elements in other labyrinthulomycetes and stramenopiles

There have been a few reports of viruses infecting labyrinthulomycetes. Takao et al. (2005, 2006) isolated a lytic ssRNA virus (SssRNAV) infecting '*Schizochytrium'* (actually an *Aurantiochytrium* strain) with a genome ~10.2kbp. Takao et al. (2007) isolated a lytic large DNA virus infecting *Sicyoidochytrium minutum* (SmDNAV), and Murakoshi et al. (2021) published its genome sequence (236,345bp with 358 predicted CDS). Takao et al. (2015) detected both types of viruses in water samples collected in Hiroshima Bay. Pollak (1979) reported an endogenous RNA virus, activated by growth in anaerobic conditions, in the thraustochytrid *Thruastochytrium aureum* (ATCC34304). We are not aware of published evidence that the ATCC MYA-1381 isolate of *A. limacinum* produces viral particles, and we have found no sequence similarity between CE1 or LE-Chr15 and SmDNAV, although we did detect genes similar to genes on SmDNAV elsewhere in the *A. limacinum* genome (not shown).

Among previous reports of NCDLV proteins in thraustochytrids, Gallot-Lavallée and Blanc (2017) detected DNA polymerase in *Aurantiochytrium limacinum* ATCC MYA-1381, plus MCP and very late transcription factor 3 (VLTF3) in *Schizochytrium aggregatum* ATCC 28209 and in *'Thraustochytrium'* LL1Fb. The *Aurantiochytrium limacinum* ATCC MYA-1381 DNA polymerase was considered a possible member of the phaeoviruses, but the others could not be placed in a known NCLDV clade. Using NCLDVOG, we confirmed the presence of these genes in the *Aurantiochytrium limacinum* ATCC MYA-1381 and *Schizochytrium aggregatum* ATCC 28209 assemblies. The DNA polymerase (NCVOG0038, blast done with AET73919) is Aurli_113265, which is one of the few predicted proteins on Scaffold_34 to have any annotation. Just downstream is an RNA polymerase large subunit (Aurli_161734 replacing the short gene model Aurli_58941), (NCVOG0274, blast done with AEQ60694 and YP_003987013). Additionally, Messyasz et al. (2020)identified a virus-like topoisomerase II on Scaffold_34 (Aurli_142950), another of the few annotated genes on this scaffold.

Among other stramenopiles, an endogenized giant viral genome is physically integrated into a nuclear chromosome in the oomycete *Phytophthora parasitica* (in a gene-sparse region rich in transposable elements) and is mainly not expressed (Hannat et al. 2021). Two potentially similar virus-like scaffolds have also been identified in *Hyphochytrium catenoides*, another deep-branching fungus-like stramenopile lineage (Leonard et al. 2018). NCLDVs, both lytic and persistent, are common among the phototrophic stramenopiles, including the phaeovirus-1 (EsV-1) domesticated in the brown alga *Ectocarpus siliculosus* (Delaroque and Boland 2008). Beyond stramenopiles, giant endogenous viral elements (EVEs) have been found in the nuclear genomes of diverse chlorophyte algae (Moniruzzaman, Weinheimer, et al. 2020), and a 400 Kb NCLDV-like scaffold has been identified in the freshwater animal *Hydra magnipapillata* (Filée 2014).

#### Potential virophage and other viral DNA in *A. limacinum* genome

A gene annotated as a regulator of chromatin condensation (RCC1) repeat-containing protein (Aurli1_13512; IPR009091) is found within a ~20kb region that is present on at least four different chromosomes (stars in **Fig. 1**). On the basis of this gene, VirSorter2 (Guo et al. 2021) classifies part of this region (scaffold_71 in the 454 assembly) as a virophage (family Lavidaviridae, viruses dependent on co-infection with large *Varidnaviria* viruses) (Bekliz, Colson, and La Scola 2016; Fischer 2021) (**Table S9**). VirSorter also identified a region on Chr17 in the Nanopore assembly as a potential dsDNA phage. Whether the putative virophage sequences detected represent parasites of mirusviruses, independent viruses infecting *A. limacinum,* or simply spurious sequence similarity will require future exploration.

### Table S1. Nanopore assembly information.

Note: this table is located in Additional File 2. Additional File 2

### Table S2. 454 Assembly information.

Note: this table is located in Additional File 2. Additional File 2

### Table S3: RepeatMasker Output

The main difference between the two assemblies in RepeatMasker is a 10-fold greater content of small RNA (pseudo) genes (0.54%) in the Nanopore, reflecting mainly the presence of rRNA genes in the Nanopore assembly contrasting with their absence in the 454 assembly.

#### 454 Assembly

```
==================================================
file name: Aurli1_sort
sequences:           181
total length:   60926267 bp  (59990477 bp excl N/X-runs)
GC level:         45.17 %
bases masked:    2213278 bp ( 3.63 %)
==================================================
               number of      length   percentage
               elements*    occupied  of sequence
--------------------------------------------------
SINEs:               48         3161 bp    0.01 %
      ALUs            0            0 bp    0.00 %
      MIRs           16         1046 bp    0.00 %
LINEs:              391        25965 bp    0.04 %
      LINE1          14          711 bp    0.00 %
      LINE2         106         6593 bp    0.01 %
      L3/CR1        128         8793 bp    0.01 %
LTR elements:        17         1010 bp    0.00 %
      ERVL            1           49 bp    0.00 %
      ERVL-MaLRs      1           43 bp    0.00 %
      ERV_classI      6          291 bp    0.00 %
      ERV_classII     1           55 bp    0.00 %
DNA elements:       129         8458 bp    0.01 %
     hAT-Charlie     13          901 bp    0.00 %
     TcMar-Tigger    14          869 bp    0.00 %
Unclassified:         3          210 bp    0.00 %
Total interspersed repeats:    38804 bp    0.06 %

Small RNA:          343        28830 bp    0.05 %

Satellites:           2          229 bp    0.00 %
Simple repeats:   42255      1923408 bp    3.16 %
Low complexity:    4421       222412 bp    0.37 %
==================================================

* most repeats fragmented by insertions or deletions
  have been counted as one element
The query species was assumed to be homo sapiens
RepeatMasker version 4.1.2-p1 , default mode
run with rmblastn version 2.2.27+
FamDB: CONS-Dfam_3.3
```

### Nanopore Assembly

```
==================================================
file name: Canu_all_reorg_27.fas
sequences:            28
total length:   62001017 bp  (62001017 bp excl N/X-runs)
GC level:         45.11 %
bases masked:    2537096 bp ( 4.09 %)
==================================================
               number of      length   percentage
               elements*    occupied  of sequence
--------------------------------------------------
SINEs:               47         2919 bp    0.00 %
      ALUs            0            0 bp    0.00 %
      MIRs           13          824 bp    0.00 %
LINEs:              390        25769 bp    0.04 %
      LINE1          21         1186 bp    0.00 %
      LINE2         100         6229 bp    0.01 %
      L3/CR1        135         8993 bp    0.01 %
LTR elements:        16          920 bp    0.00 %
      ERVL            1           49 bp    0.00 %
      ERVL-MaLRs      2           96 bp    0.00 %
      ERV_classI      7          372 bp    0.00 %
      ERV_classII     0            0 bp    0.00 %
DNA elements:       125         8431 bp    0.01 %
     hAT-Charlie     12          880 bp    0.00 %
     TcMar-Tigger    13          815 bp    0.00 %
Unclassified:         2          122 bp    0.00 %
Total interspersed repeats:    38161 bp    0.06 %

Small RNA:          727       344359 bp    0.56 %

Satellites:           2          229 bp    0.00 %
Simple repeats:   41818      1931247 bp    3.11 %
Low complexity:    4350       223559 bp    0.36 %
==================================================

* most repeats fragmented by insertions or deletions
  have been counted as one element
The query species was assumed to be homo sapiens
RepeatMasker version 4.1.2-p1 , default mode
run with rmblastn version 2.2.27+
FamDB: CONS-Dfam_3.3
```

### Table S4. BUSCO 5.3 results from dataset eukaryota_odb10 on the Nanopore and 454 genome assemblies.

Most of the difference in complete BUSCOs is due to more Fragmented BUSCOs in the Nanopore assembly, likely reflecting lower sequence accuracy of the long-read technology. In BUSCO analyses of 10 labyrinthulomycete genomes and transcriptomes from 7 species (Collier, Rest et al. in preparation), the *Aurantiochytrium limacinum* 454 assembly is assessed as the most complete, and 11 BUSCOs are missing from all (some replaced by bacteria HGT, some missed by BUSCO because of gene fusions), suggesting that their absence reflects features of labyrinthulomycetes evolution rather than low-quality genome assemblies.

| BUSCOs (n:255) | Nanopore | % | 454 | % |
|---|---|---|---|---|
| Complete (C) | 187 | 73.4% | 222 | 87.1% |
| Complete and single-copy (S) | 184 | 72.2% | 221 | 86.7% |
| Complete and duplicated (D) | 3 | 1.2% | 1 | 0.4% |
| Fragmented (F) | 37 | 14.5% | 11 | 4.3% |
| Missing (M) | 31 | 12.1% | 22 | 8.6% |

### Table S5. LOng REpeated TElomere And Rrna Spacers (LORE-TEARS) statistics

| descriptor | | group 0 | group 1 | group 2 | group 3 | group 5 | Others | Telomeres |
|---|---|---|---|---|---|---|---|---|
| # of repeat units | mean | 9.3 | 3.8 | 5.7 | 2.9 | 5.4 | | 102.4393 |
| | median | 8.6 | 3.9 | 5.2 | 2.9 | 4.6 | | 100.2 |
| | min | 1.8 | 2.5 | 1.8 | 1.9 | 1.8 | | 48.8 |
| | max | 20.6 | 5 | 15.9 | 4 | 20.4 | | 159 |
| length of repeat unit | mean | 391 | 407 | 365 | 529 | 434 | | |
| | median | 385 | 407 | 366 | 530 | 430 | | |
| | min | 326 | 398 | 347 | 515 | 408 | | |
| | max | 627 | 413 | 370 | 533 | 481 | | |
| # of elements (Chr 1-26) | | 94 | 44 | 75 | 19 | 32 | 88 | |
| # with G-quadruplex | | 15 | 0 | 0 | 3 | 1 | 14 | |
| location between | | 5S and 18S | 28S and telomeres | 28S and telomeres | group 1 and group 2 | two 5S rRNAs (typically) | various | |

### Table S6. G-quadruplexes detected within 28S rDNA genes

Note: this table is located in Additional File 2. Additional File 2

We predicted G-quadruplexes and found them to be located inside and just downstream of many of the rRNA clusters (**Fig. 2**). For 72 of the 73 28S subtelomeric rRNA genes, a G-quadruplex is predicted within the rRNA coding region, about 3/5 of the way into the gene. For 52 of the 28S rRNAs, a G-quadruplex is predicted about 60 bp downstream of the predicted last nucleotide of the mature RNA. We also found that 32% of Group 0 LORE-TEARS elements contain a G-quadruplex (**Table S5**). G-quadruplexes can have important regulatory functions, including at chromosome ends (Juranek and Paeschke 2012; Paeschke et al. 2008; Biffi, Tannahill, and Balasubramanian 2012; Wang et al. 2012), and their prediction in the LORE-TEARS repeats and downstream of the rRNA clusters is consistent with a regulatory function for these elements at the chromosome ends of *A. limacinum*.

### Table S7: Viral content detected in *A. limacinum* predicted proteins by HMM and BLAST.

This table includes all reciprocal best BLASTp hits between proteins on CE1 and Chr15.
Note: this table is located in Additional File 2. Additional File 2

### Table S8: Viral (NCLDV) sequences identified by ViralRecall

Region-scale parameters: -m 75 -g 4 -w 175
All results with scores > 0 are shown. For contig/scaffold (-c) level results (bottom three rows), only contigs and scaffolds >290kb are shown.

| **Assemb-ly** | **Scale** | **replicon** (chromosome) [note] | **start coord** | **end coord** | **region or contig length** | **score** | **num viral hits** | **num ORFs** | **num pfam hits** | **markers** |
|---|---|---|---|---|---|---|---|---|---|---|
| 454 | region | scaffold_35(Chr15) | 45406 | 294615 | 249209 | 0.620 | 34 | 207 | | RNR,RNAPL,PolB |
| 454 | region | rcscaffold_34(CE1) | 9316 | 297367 | 288051 | 0.205 | 31 | 228 | 37 | PolB,RNAPL |
| Nanopore | region | tig00000136(CE1) | 50971 | 298422 | 247451 | 0.183 | 31 | 305 | 34 | RNAPL |
| Nanopore | region | tig00004069(Chr15) | 111653 | 278329 | 166676 | 0.442 | 33 | 239 | | RNR,RNAPL |
| 454 | scaffold | scaffold_35(Chr15) | | | 295699 | 0.512 | 42 | 241 | 45 | PolB,RNAPL,RNR |
| 454 | scaffold | rcscaffold_34(CE1) | | | 297369 | 0.185 | 32 | 234 | 38 | PolB,RNAPL |
| Nanopore | contig | tig00000136(CE1) | | | 298423 | 0.113 | 38 | 368 | 43 | RNAPL |

### Table S9: Viral sequences identified by VirSorter

*scaffold_71 is 3888 bp and was not aligned by Mauve with the Nanopore assembly, but Blast identified similar sequences at four locations in the Nanopore assembly: Chr10, Chr18, Chr20, Chr21.
**tig00000194 is 39992bp, and the only unassembled Nanopore contig lacking both telomeric repeats and rRNA genes.

| assembly | seqname | class | score | length | hallmark | viral | cellular |
|---|---|---|---|---|---|---|---|
| 454 | scaffold_35(Chr15) | NCLDV | 0.727 | 273131 | 0 | 6 | 0 |
| 454 | rcscaffold_34(partial)(CE1) | NCLDV | 0.853 | 191208 | 1 | 6.2 | 1.2 |
| 454 | scaffold_71\|\|full* | lavidaviridae | 0.993 | 3885 | 0 | 25 | 0 |
| Nanopore | tig00000194\|\|full** | NCLDV | 0.773 | 36765 | 1 | 22.7 | 0 |
| Nanopore | tig00004069\|\|0_partial(Chr15) | dsDNAphage | 0.887 | 22662 | 1 | 3.6 | 0 |
| Nanopore | tig00004074\|\|0_partial(Chr17) | dsDNAphage | 0.9 | 22925 | 1 | 10.3 | 0 |

### Figure S1. Mauve mapping

S1A: MAUVE alignment of (top to bottom): the combined Canu assembly, wild-type assembly, *crtIBY* knockout 32 and *crtIBY* knockout 33.

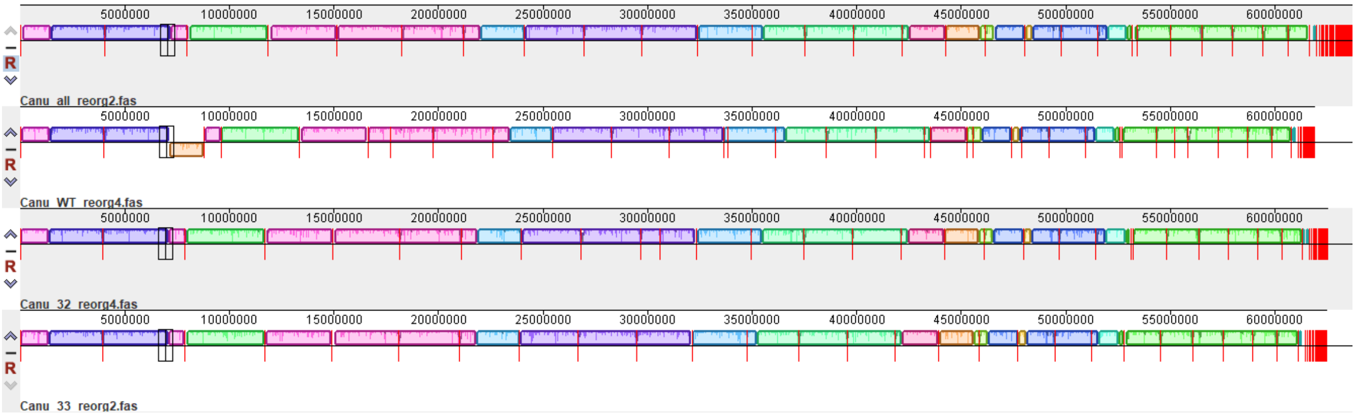


S1B: 454 assembly (bottom) mapped to the Nanopore assembly (top). Numbers immediately below each assembly are the final non-zero digits of scaffold and contig numbers. White numbers on black background are proposed chromosome numbering, from largest to smallest of the first 27 contigs in the Nanopore assembly.

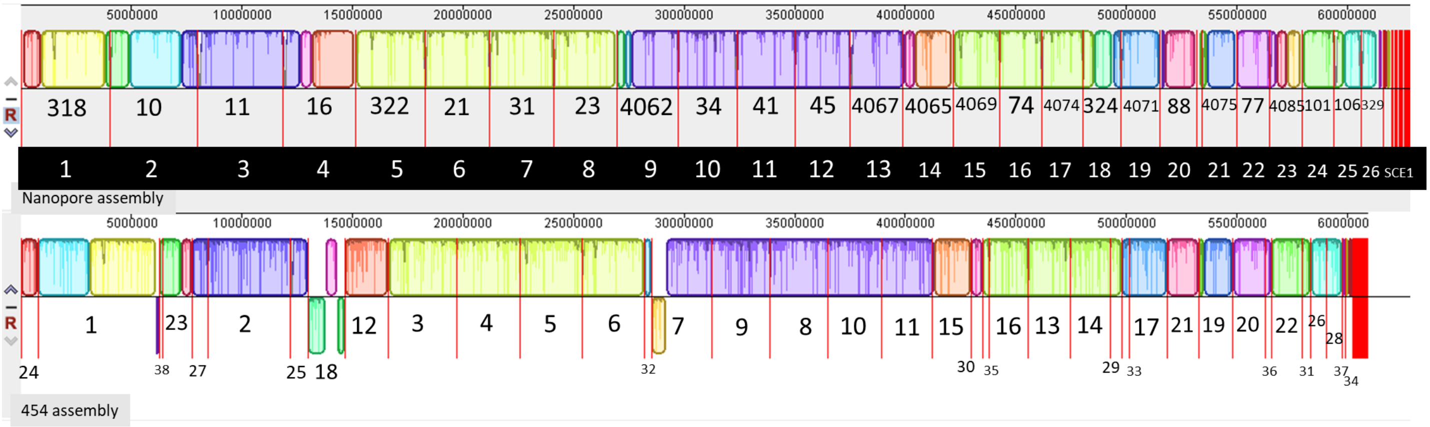

### Figure S2. Mapping between Nanopore and 454 Assemblies

#### Figure S2A.

Mapping between the Nanopore-based 26 putative linear chromosomes (and CE1) and the 454-based assembly of *Aurantiochytrium limacinum* ATCC MYA-1381. Circular Element 1 is predicted to be circular, but is displayed as linear. A scale in megabases is provided along the top of the plot (Gel and Serra 2017). Vertical red lines represent locations of predicted (Lagesen et al. 2007) rRNA gene regions, and are found almost exclusively at the ends of the linear chromosomes. Cyan boxes represent regions that did not align, via Mauve (Darling et al. 2004) with the primary (long scaffolds) of the 454 assembly. Yellow boxes represent regions that are found (via BLAST) to be present in the short scaffolds of the 454 assembly.

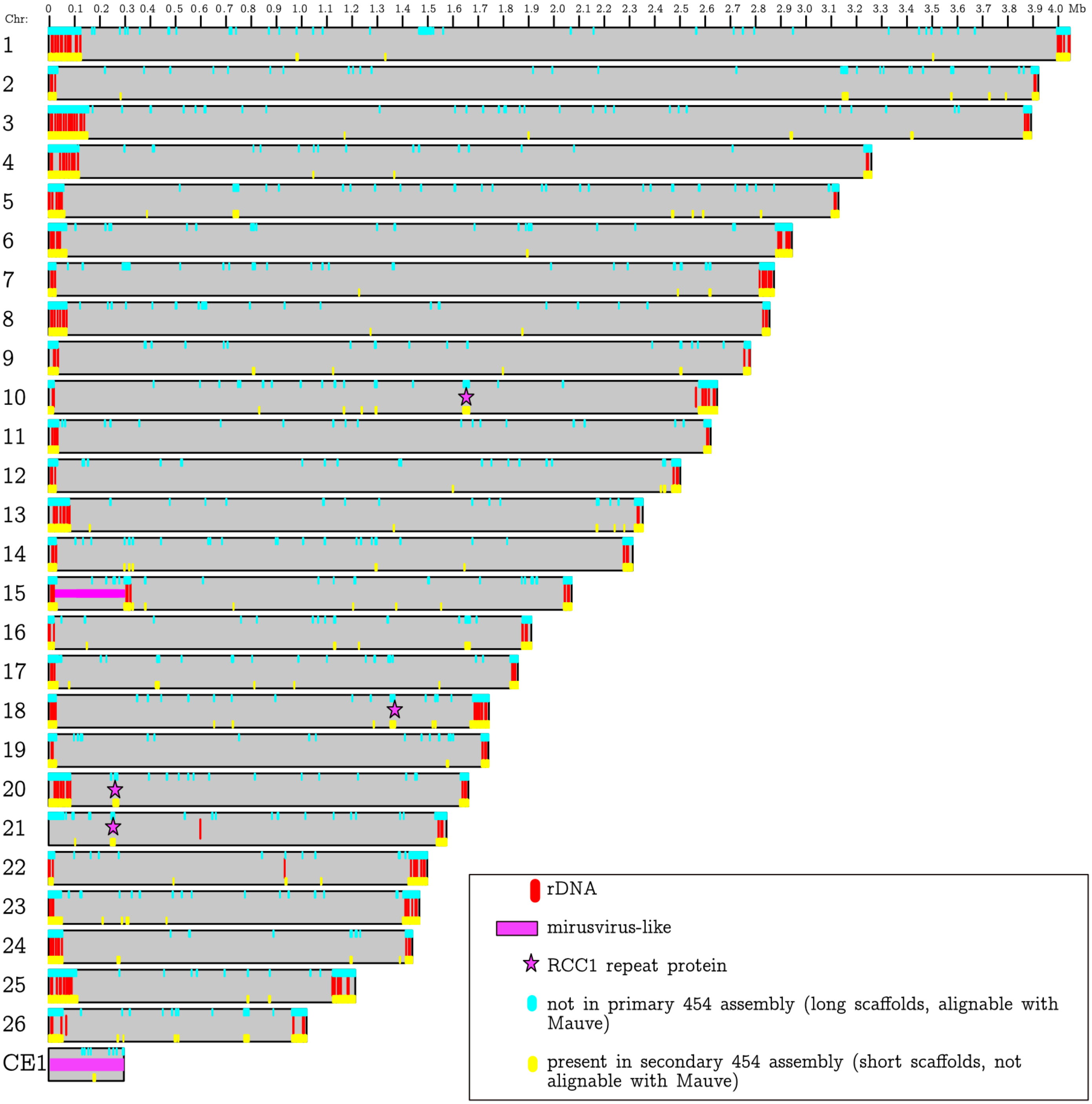

### Fig. S2B.

Focused view of the 50 Kb of both ends of the *Aurantiochytrium limacinum* chromosomes. This is identical to Fig. 2, but with regions that do not map to the primary 454 assembly (cyan), and regions that do map to the secondary 454 assembly (yellow) annotated. The primary assembly is composed of long 454 scaffolds that are alignable to the Nanopore assembly with Mauve, while the secondary assembly is composed of short scaffolds that are not alignable to the Nanopore assembly with Mauve (and were mapped with BLAST). Also indicated: Locations of predicted rRNA genes (red); telomeric repeats (purple), LORE-TEARS (orange, brown, green), G-quadruplexes (blue lines), exons (black lines), GC content (plotted along the centerline).

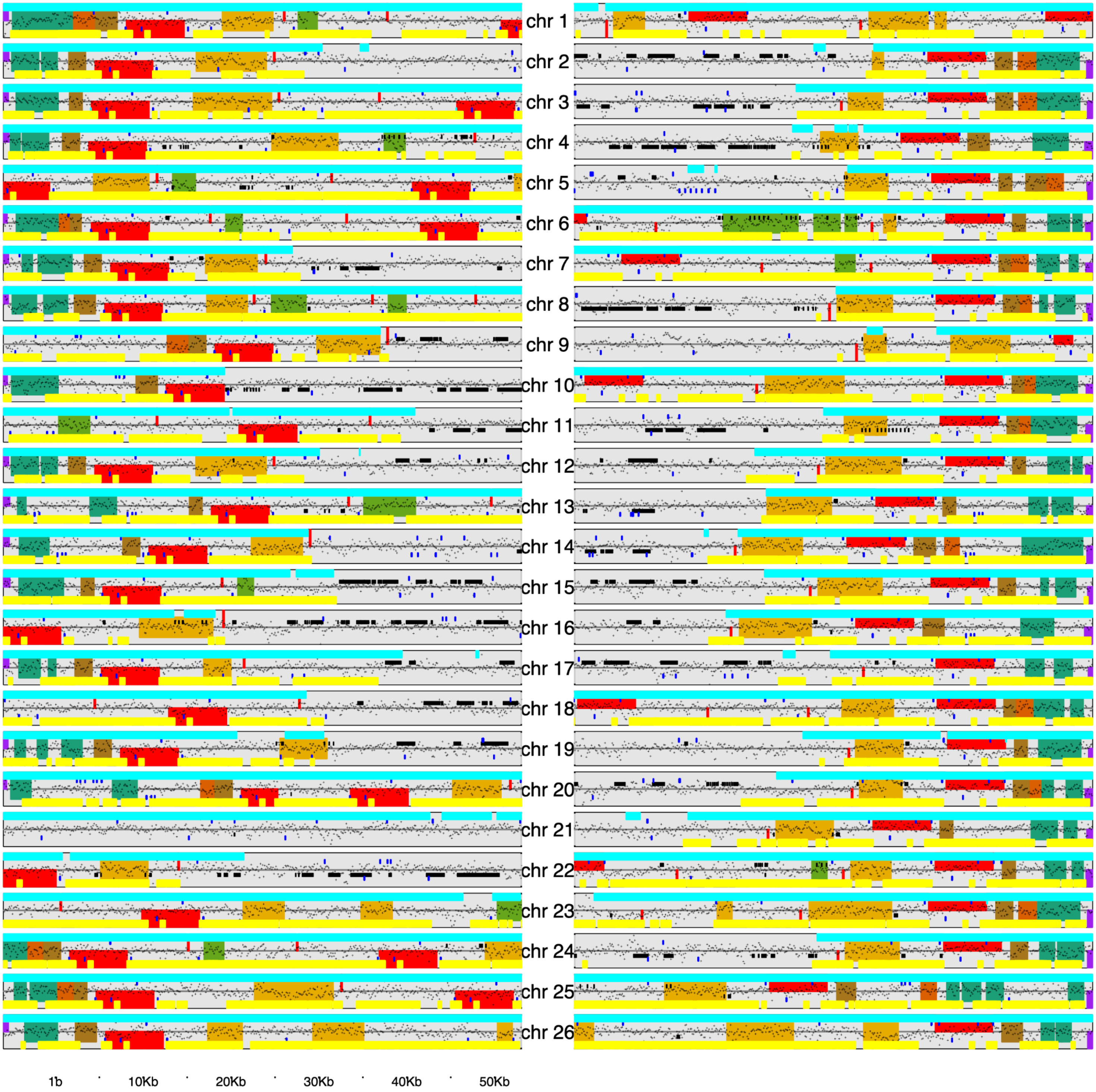

### Figure S3. Physical chromosome sizing

Pulsed-field gel electrophoresis of *Aurantiochytrium limacinum* MYA-1381 chromosomes. Chromosome-sized bands ranging in size from >3.0 to ~0.30 Mbp were apparent, consistent with the assembled genome. Some bands presumably represent more than a single chromosome. Abbreviations: Sc, *Saccharomyces cereviseae*; Aurantio, *A. limacinum*. Anbu et al. (2007) reported similar analysis for four different thraustochytrids, finding between 8 and 17 chromosomes ranging from 0.220 to 2.205 Mbp, with estimated genome sizes between 9.93 and 12.9 Mbp. We suspect these are underestimates arising from the comigration of chromosomes of similar size and larger chromosomes unresolved in the top of the gel.

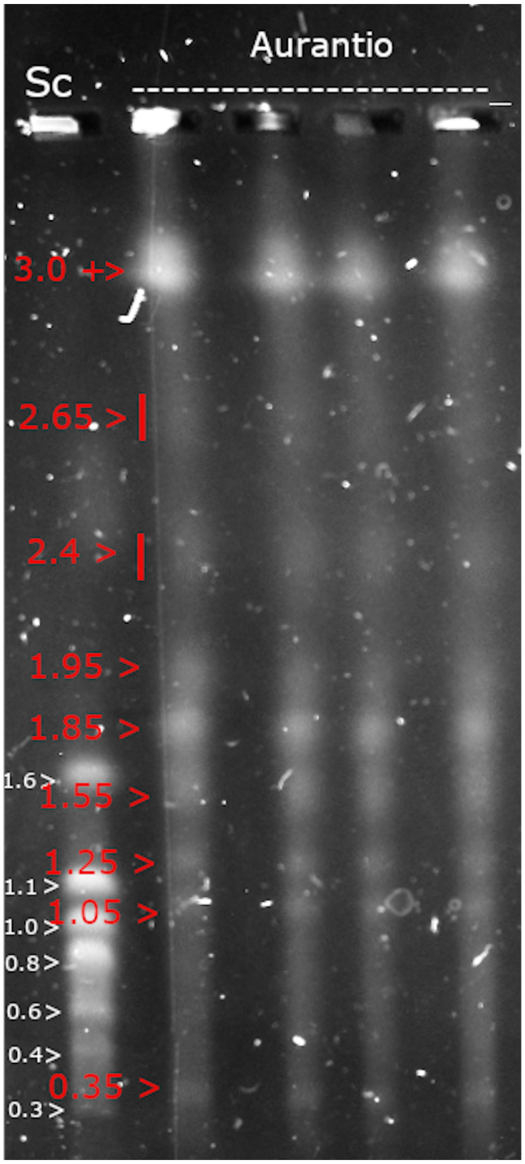

### Figure S4. ASGART comparison

Chord plot showing matching sequence regions between scaffolds in 454 (left) and contigs in Nanopore (right) -based assemblies. Scaffolds/contigs are arbitrarily colored to make visualization distinct. Chord connections indicate regions of similarity of at least 1 Kbp, as determined by ASGART (Delehelle et al. 2018). An arbitrary set of chords are colored red and blue to highlight the directional nature of the repeats at the scaffold/contig ends. A blank space in both assemblies represents the CE1 scaffold/contig, which has no matching sequence regions on other scaffolds/contigs.

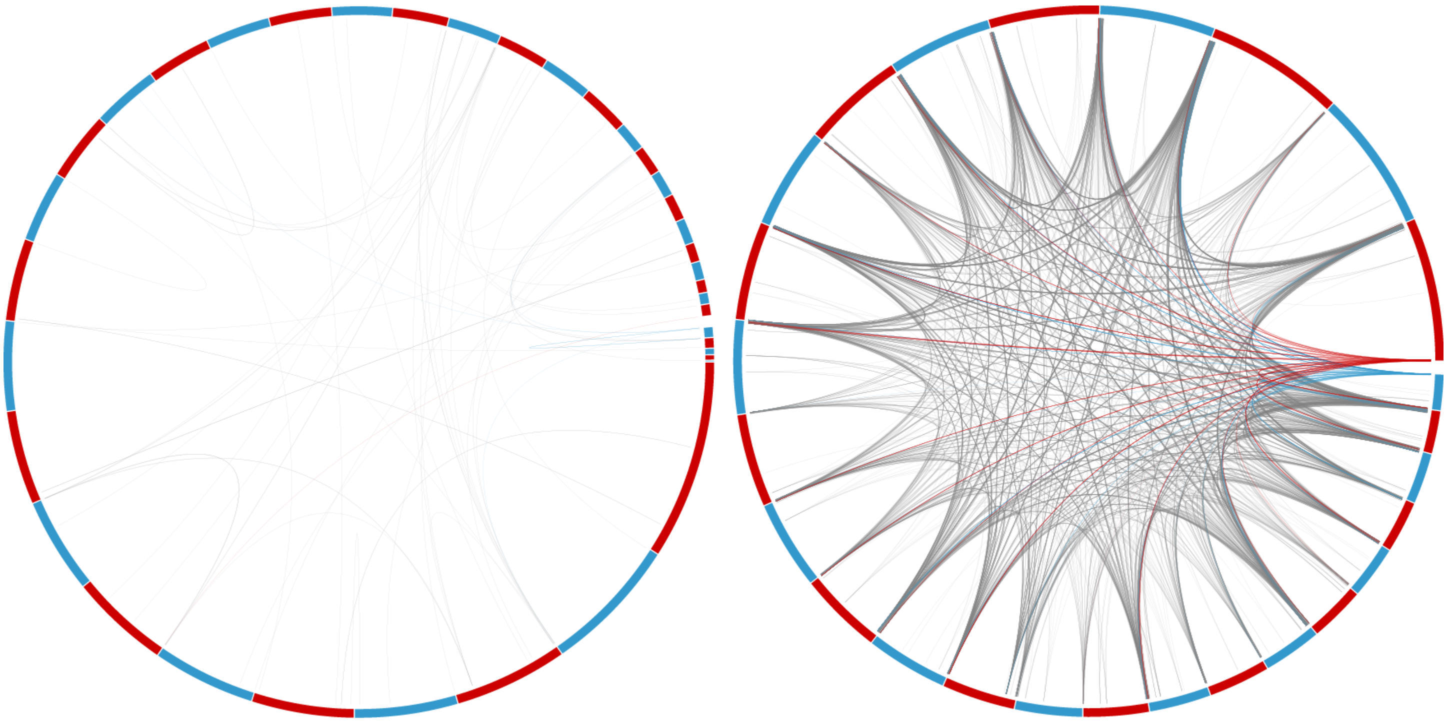

### Figure S5. Nanopore tig00000136 assembly and 454 scaffold_34 assembly dotplots

The shortest of the putative chromosomal elements in *A. limacinum* (excluding the mitochondrial genome) was represented by tig00000136 in the Nanopore assembly and Scaffold_34 in the 454 assembly (357 and 297 kb, respectively). The Canu-based Nanopore assembly predicts that this element is circular; BLAST comparisons showed that it contains ~60 Kbp of nearly identical sequence at each end. Scaffold_34 (from the 454 assembly) corresponds to a linear version of the same element linearized at a different point. Circularization of tig00000136 by collapsing the ends resulted in a ~298 Kbp element, which we consider to be the 27th genomic element of *A limacinum,* dubbed CE1 (circular element 1).

Below, BLASTn of Nanopore Chromosome 27 (tig00000136) vs. itself (top) and vs 454 scaffold_34 (bottom). The Canu assembly repeats ~60kb at each end of the break; this collapses once circularized. The unique length of the scaffold is 298424 bp.

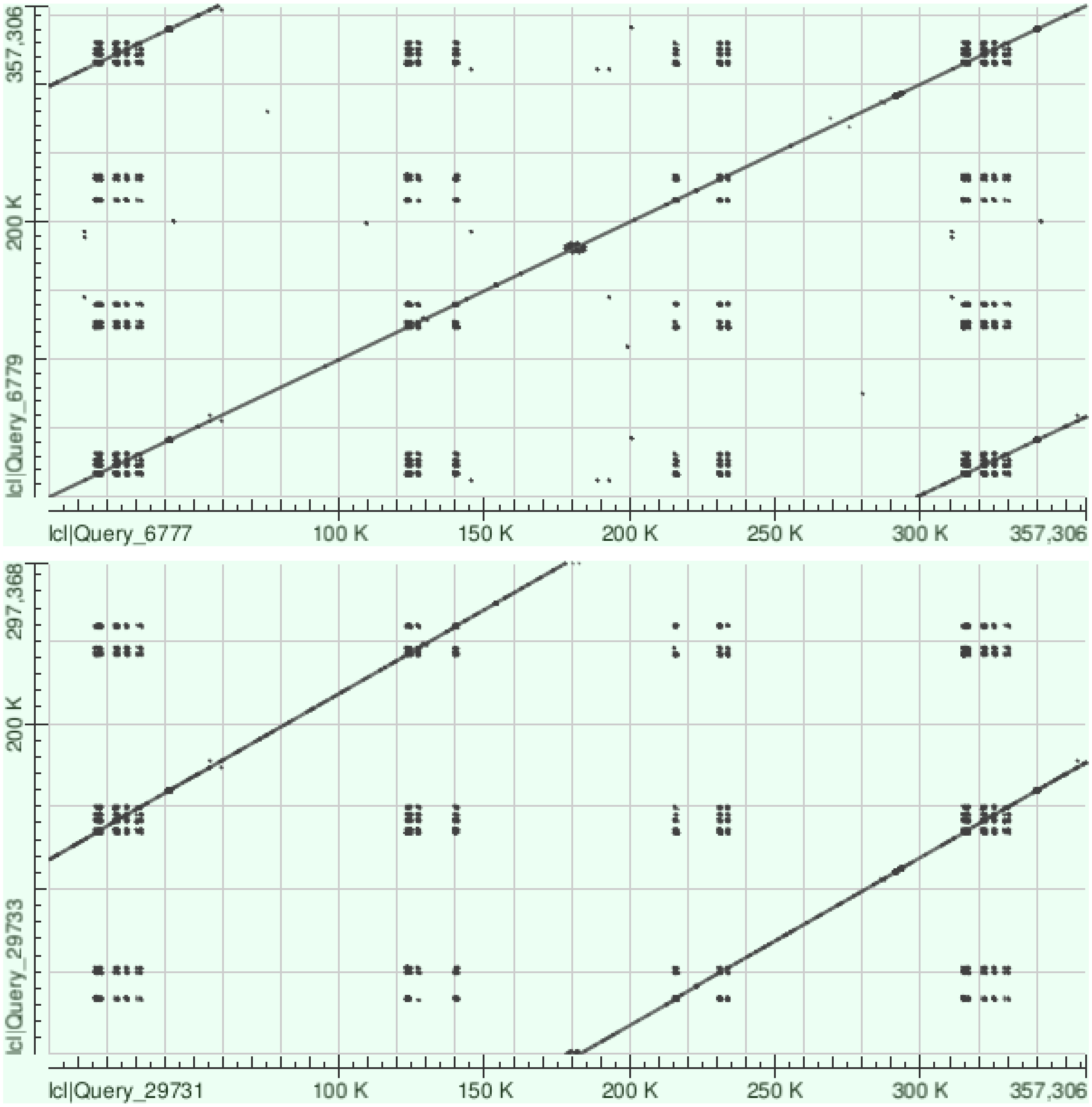

### Figure S6. Proportion of genes missing annotations across scaffolds

Proportion of genes on a scaffold that lack at least one annotation for each type (GO, IPR, KEGG, KOG), or for all four types, for SigP, or that lack at least one intron. Red points represent proportions for scaffold_34 (CE1).

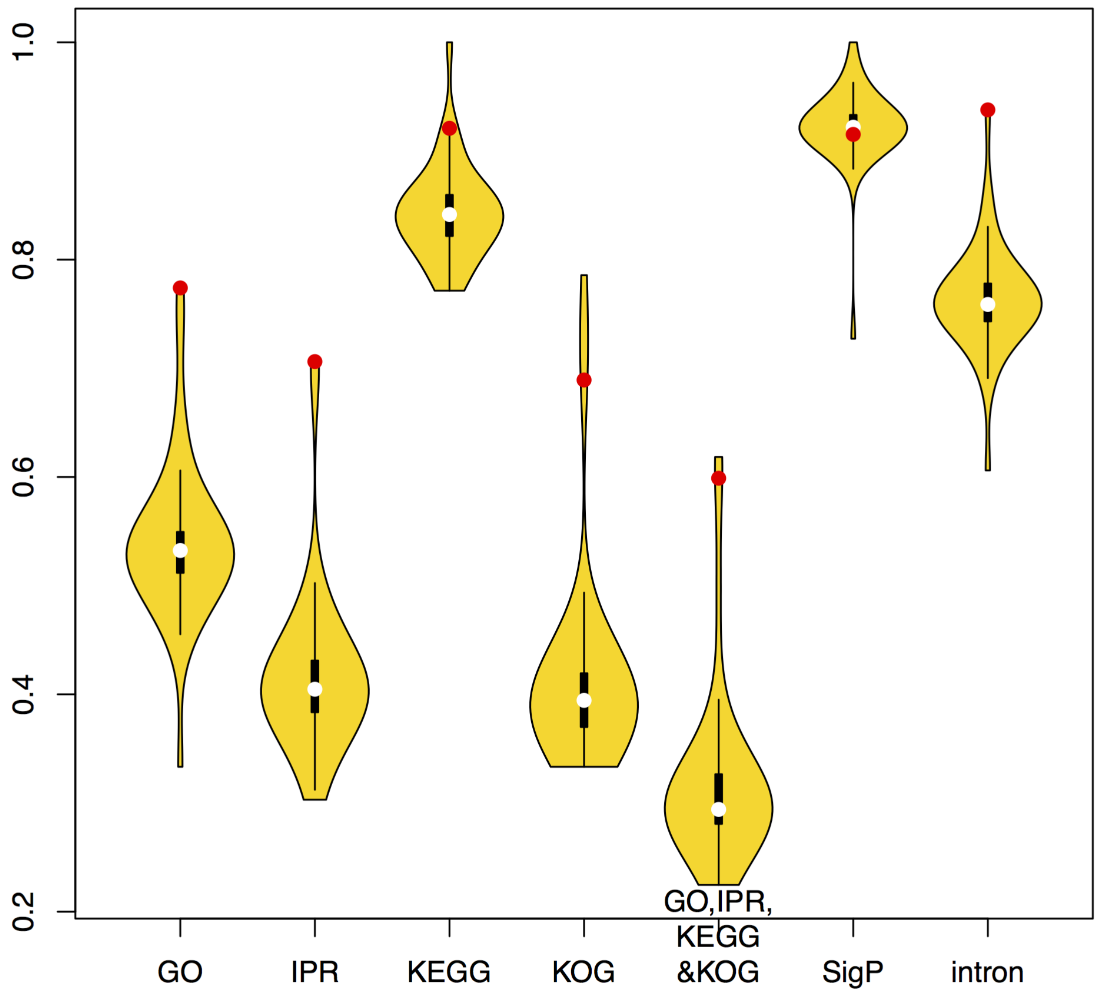

### Figure S7. Phylogenetic trees of select proteins encoded on CE1 and the LE-Chr15 viral-like element

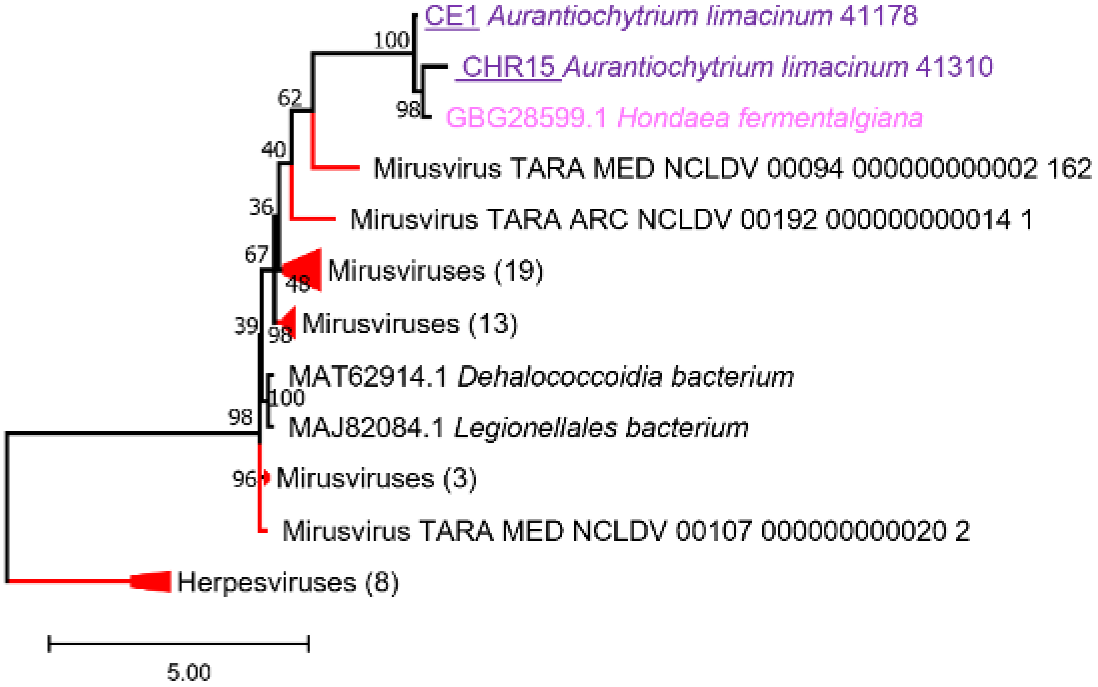


#### Fig. S7A. Phylogeny of terminases rooted with herpesviruses homologs.

*Aurantiochytrium limacinum* sequences are in purple; the chromosome on which the sequence resides is provided in the OTU name. *Hondaea fermentalgiana* sequences are in pink, viral sequences are in red, and prokaryotic sequences are in black. The scale bar indicates the inferred number of amino acid substitutions per site. The sequences were aligned with MAFFT-linsi prior to phylogenetic reconstruction.

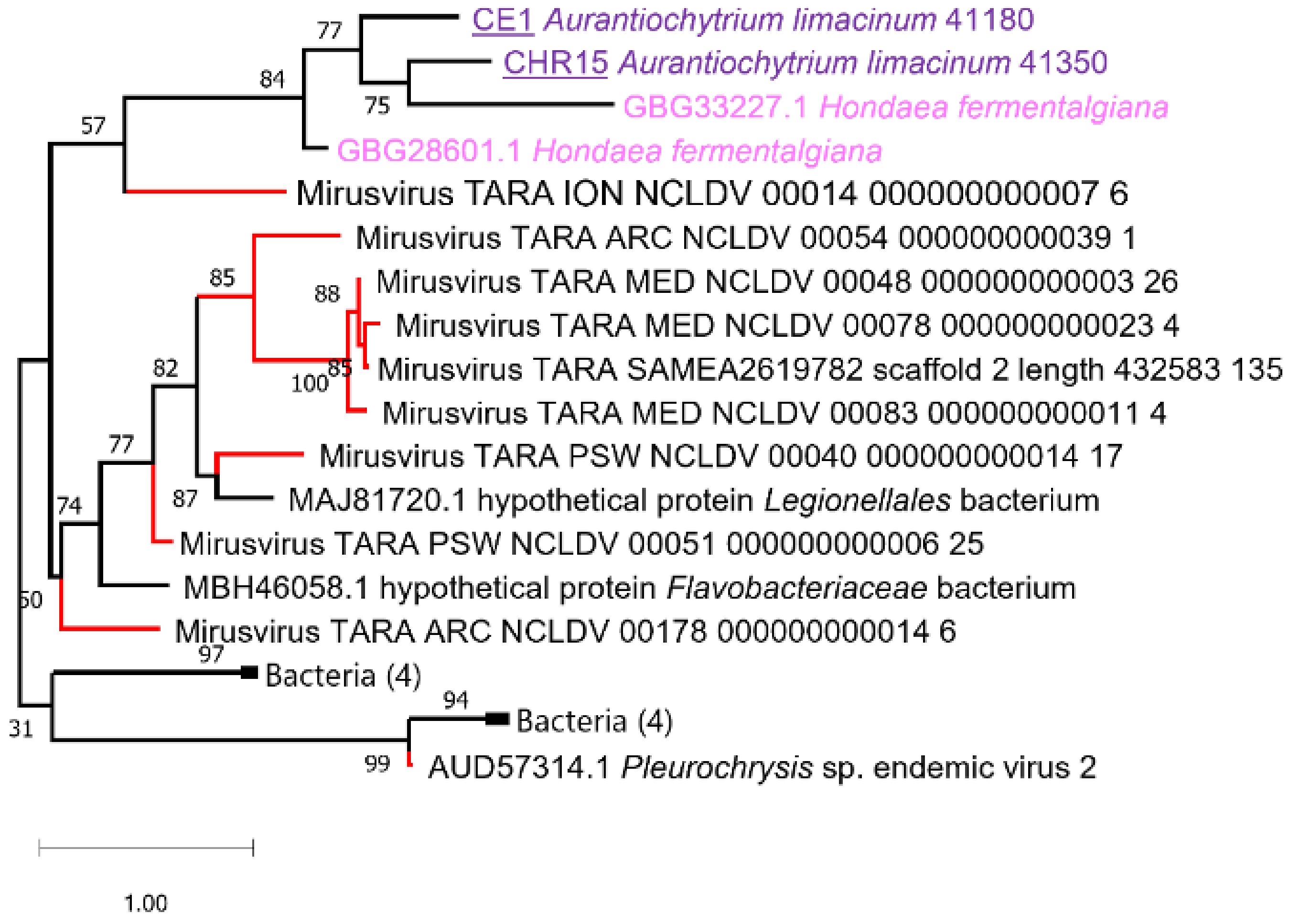


#### Fig. S7B. Phylogeny of Holliday junction resolvases.

*Aurantiochytrium limacinum* sequences are in purple; the chromosome on which the sequence resides is provided in the OTU name. *Hondaea fermentalgiana* sequences are in pink, viral sequences are in red, and prokaryotic sequences are in black. The scale bar indicates the inferred number of amino acid substitutions per site. The sequences were aligned with MAFFT-linsi prior to phylogenetic reconstruction.

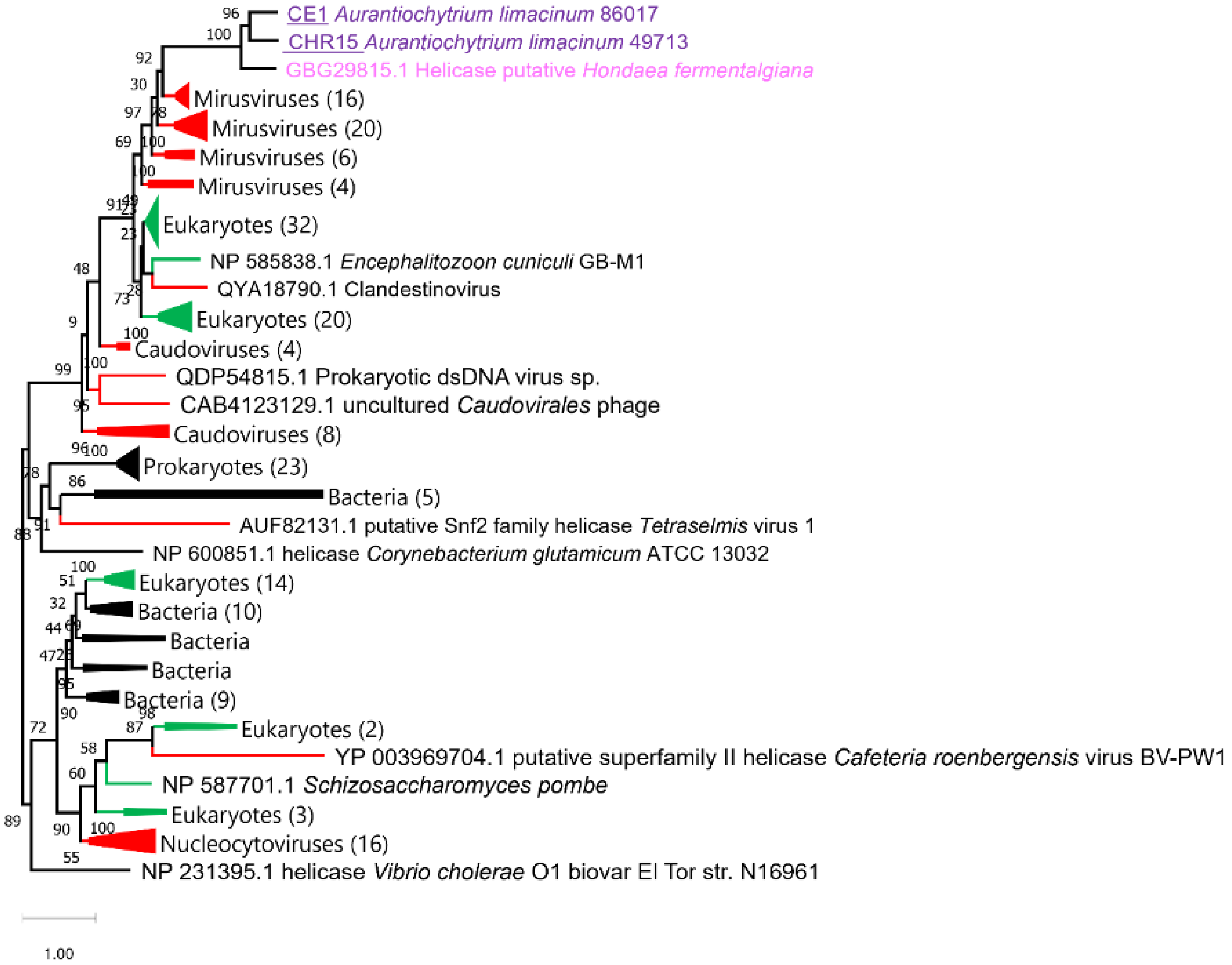


### Fig. S7C. Phylogeny of superfamily 2 helicases.

*Aurantiochytrium limacinum* sequences are in purple; the chromosome on which the sequence resides is provided in the OTU name. *Hondaea fermentalgiana* sequences are in pink, viral sequences are in red, eukaryotic sequences are in green, and prokaryotic sequences are in black. The scale bar indicates the inferred number of amino acid substitutions per site. The sequences were aligned with MAFFT and columns with less than 10% gaps were retained for phylogenetic reconstruction.

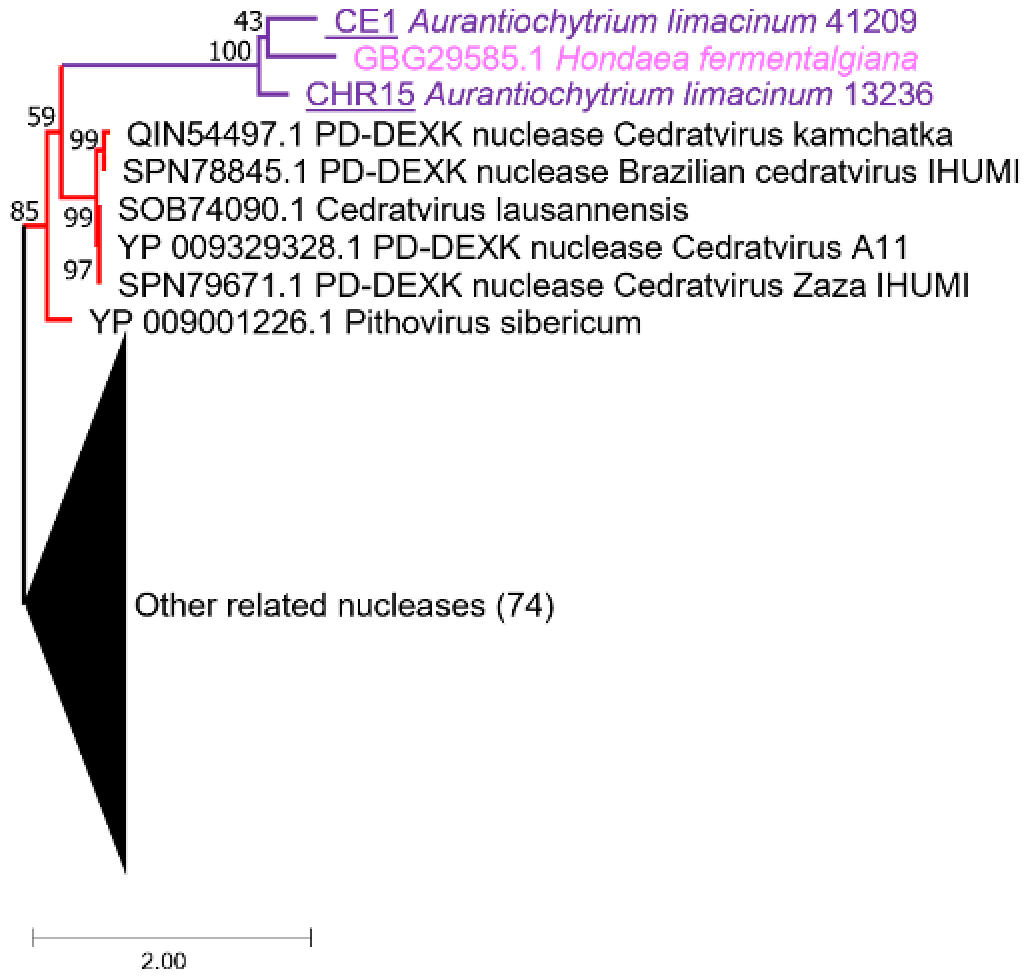


### Fig. S7D. Phylogeny of PD-DEXK nucleases.

*Aurantiochytrium limacinum* sequences are in purple; the chromosome on which the sequence resides is provided in the OTU name. *Hondaea fermentalgiana* sequences are in pink, viral sequences are in red, and prokaryotic sequences are in black. The scale bar indicates the inferred number of amino acid substitutions per site.

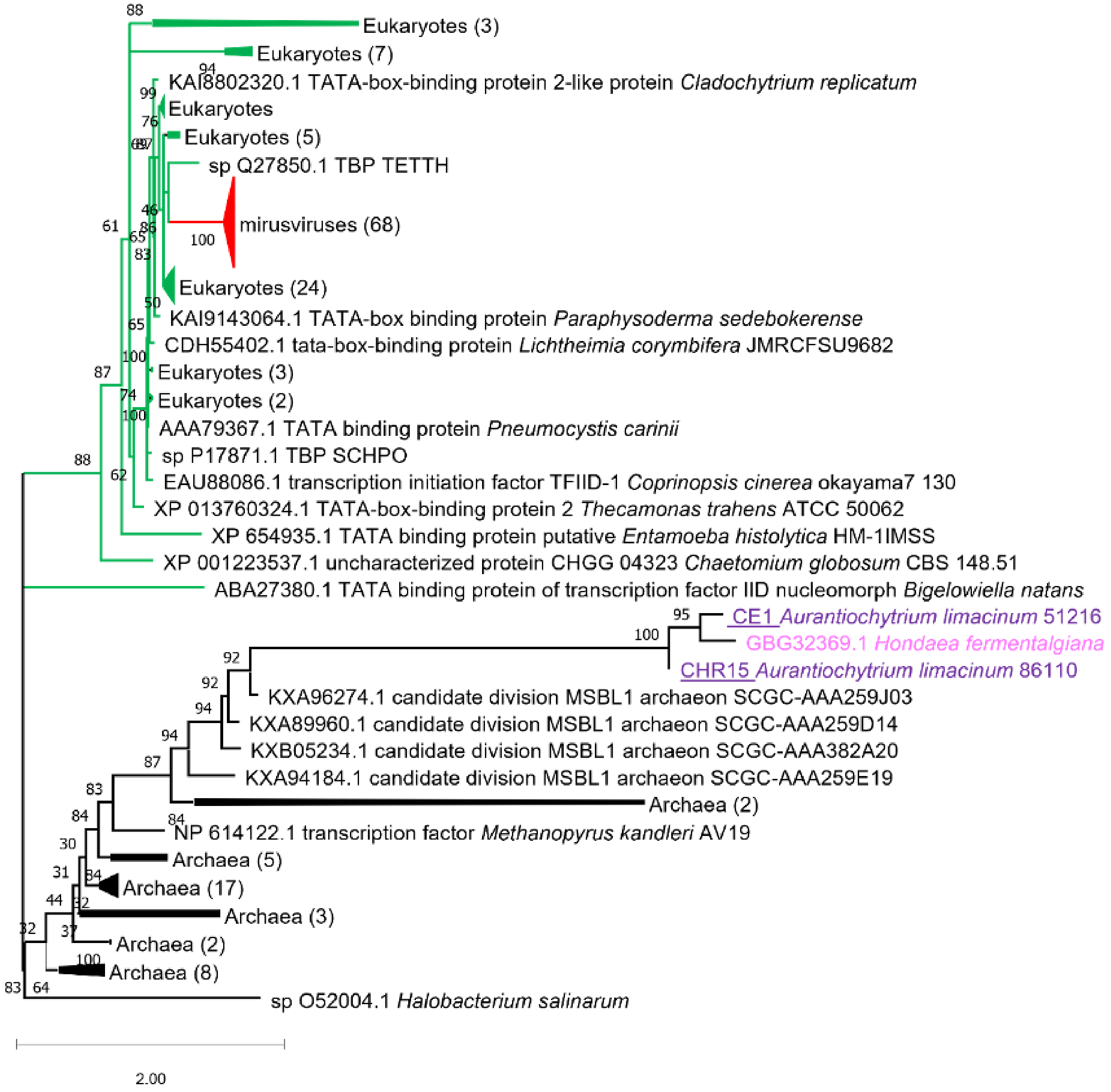


### Fig. S7E. Phylogeny of TATA box-binding proteins.

*Aurantiochytrium limacinum* sequences are in purple; the chromosome on which the sequence resides is provided in the OTU name. *Hondaea fermentalgiana* sequences are in pink, viral sequences are in red, eukaryotic sequences are in green, and prokaryotic sequences are in black. The scale bar indicates the inferred number of amino acid substitutions per site.

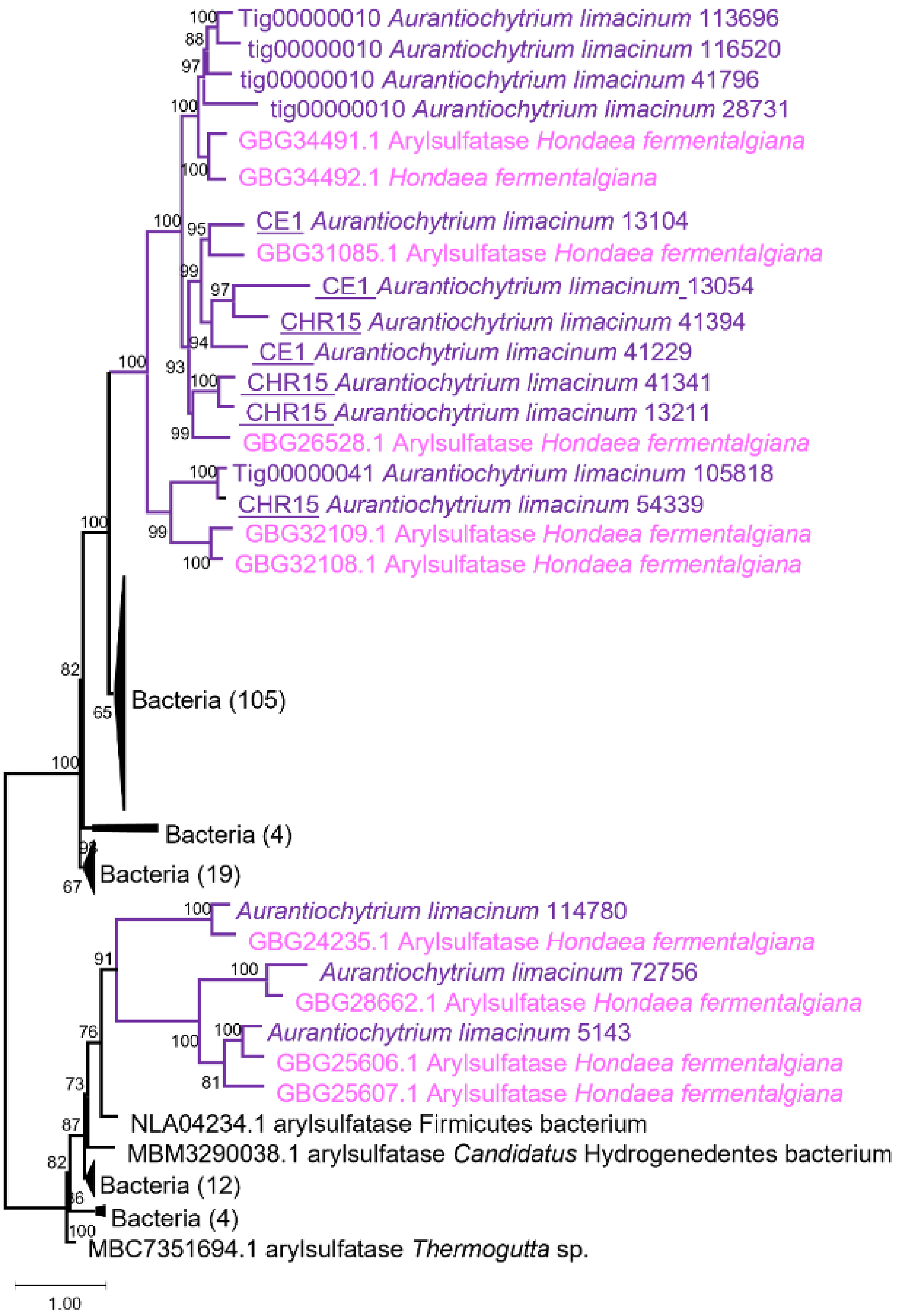


### Fig. S7F. Phylogeny of arylsufatases.

*Aurantiochytrium limacinum* sequences are in purple; the chromosome on which the sequence resides is provided in the OTU name. *Hondaea fermentalgiana* sequences are in pink, viral sequences are in red, and prokaryotic sequences are in black. The scale bar indicates the inferred number of amino acid substitutions per site.

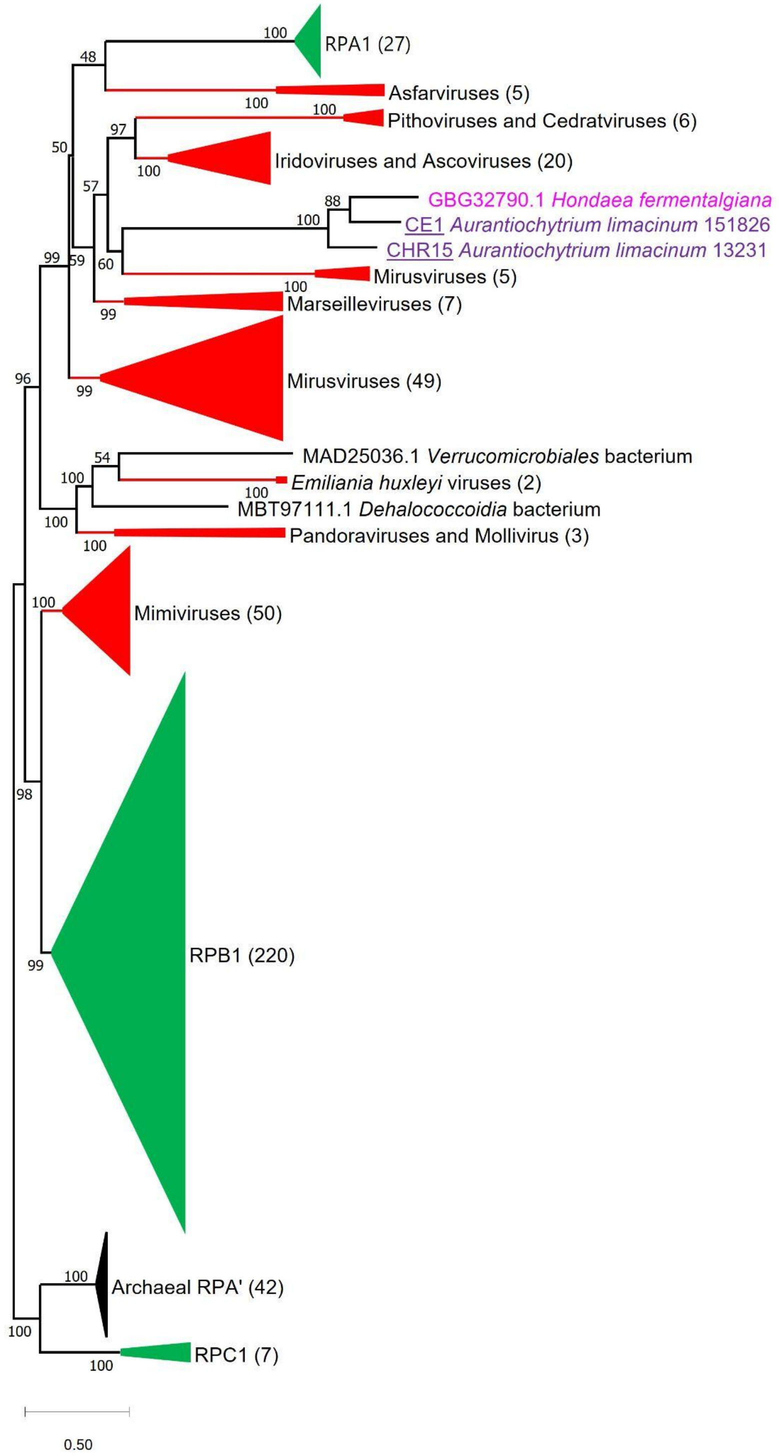


### Fig. S7G. Phylogeny of RNAPol N-terminal subunit.

*Aurantiochytrium limacinum* sequences are in purple; the chromosome on which the sequence resides is provided in the OTU name. *Hondaea fermentalgiana* sequences are in pink, viral sequences are in red, eukaryotic sequences are in green, and prokaryotic sequences are in black. The scale bar indicates the inferred number of amino acid substitutions per site.

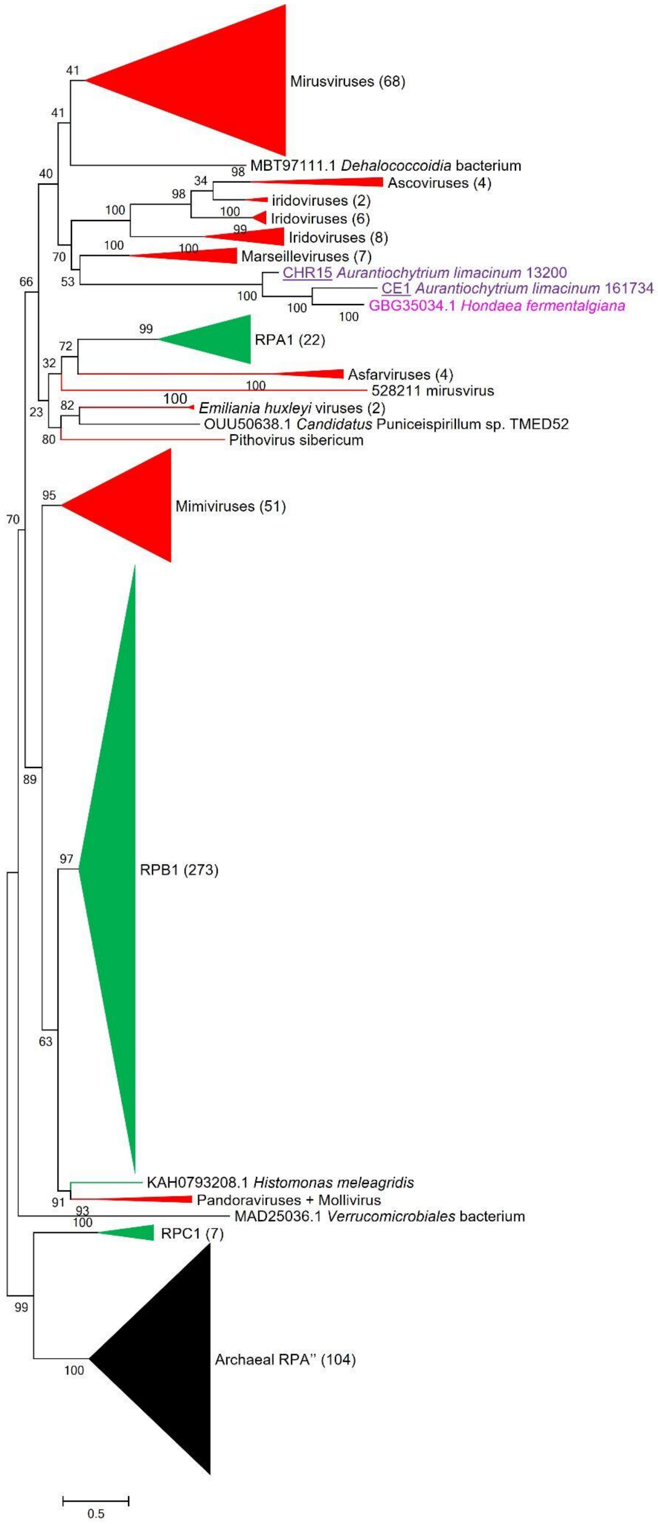


### Fig. S7H. Phylogeny of RNAPol C-terminal subunit.

*Aurantiochytrium limacinum* sequences are in purple; the chromosome on which the sequence resides is provided in the OTU name. *Hondaea fermentalgiana* sequences are in pink, viral sequences are in red, eukaryotic sequences are in green, and prokaryotic sequences are in black. The scale bar indicates the inferred number of amino acid substitutions per site.

### Figure S8. tBLASTx comparison of viral regions on CE1 and chromosome 15 (LE-Chr15)

tBLASTx based comparison of rscaffold_34 (CE1; query) and scaffold_35 (chromosome 15; subject)

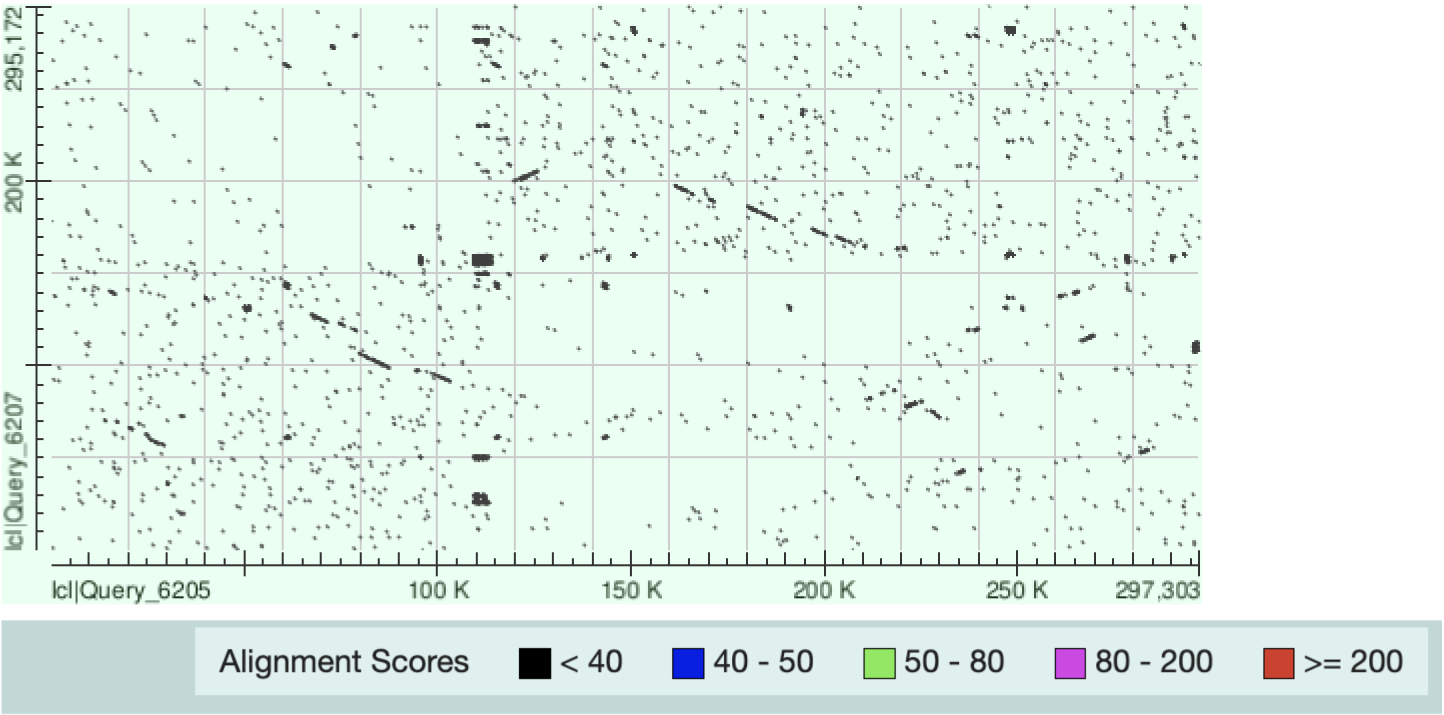


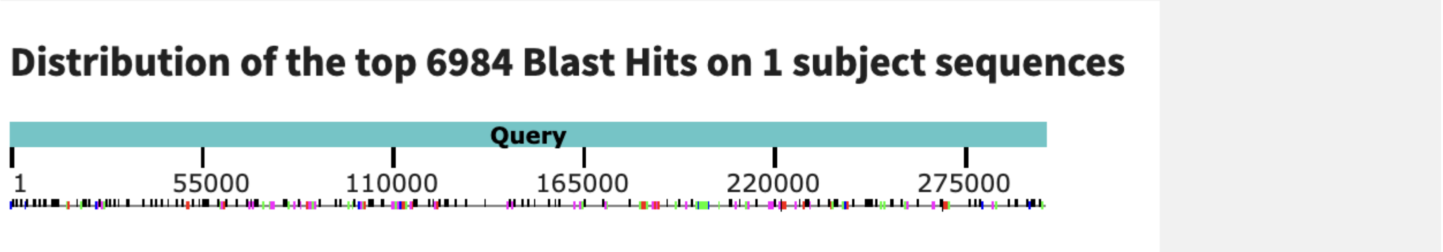


| Max Score | Total Score | Query Cover | E value | Per. Ident |
|---|---|---|---|---|
| 777 | 4.038e+05 | 43% | 0.0 | 64.15% |

Archer, John, Greg Baillie, Simon J. Watson, Paul Kellam, Andrew Rambaut, and David L. Robertson. 2012. “Analysis of High-Depth Sequence Data for Studying Viral Diversity: A Comparison of next Generation Sequencing Platforms Using Segminator II.” *BMC Bioinformatics* 13 (March): 47.

Aylward, Frank O., and Mohammad Moniruzzaman. 2021. “ViralRecall-A Flexible Command-Line Tool for the Detection of Giant Virus Signatures in ’Omic Data.” *Viruses* 13 (2): 150.

Aylward, Frank O., Mohammad Moniruzzaman, Anh D. Ha, and Eugene V. Koonin. 2021. “A Phylogenomic Framework for Charting the Diversity and Evolution of Giant Viruses.” *PLoS Biology* 19 (10): e3001430.

Bekliz, Meriem, Philippe Colson, and Bernard La Scola. 2016. “The Expanding Family of Virophages.” *Viruses* 8 (11). https://doi.org/10.3390/v8110317.

Biffi, Giulia, David Tannahill, and Shankar Balasubramanian. 2012. “An Intramolecular G-Quadruplex Structure Is Required for Binding of Telomeric Repeat-Containing RNA to the Telomeric Protein TRF2.” *Journal of the American Chemical Society* 134 (29): 11974–76.

Darling, Aaron C. E., Bob Mau, Frederick R. Blattner, and Nicole T. Perna. 2004. “Mauve: Multiple Alignment of Conserved Genomic Sequence with Rearrangements.” *Genome Research* 14 (7): 1394–1403.

Delaroque, Nicolas, and Wilhelm Boland. 2008. “The Genome of the Brown Alga Ectocarpus Siliculosus Contains a Series of Viral DNA Pieces, Suggesting an Ancient Association with Large dsDNA Viruses.” *BMC Evolutionary Biology* 8 (April): 110.

Delehelle, Franklin, Sylvain Cussat-Blanc, Jean-Marc Alliot, Hervé Luga, and Patricia Balaresque. 2018. “ASGART: Fast and Parallel Genome Scale Segmental Duplications Mapping.” *Bioinformatics* 34 (16): 2708–14.

Ewing, B., and P. Green. 1998. “Base-Calling of Automated Sequencer Traces Using Phred. II. Error Probabilities.” *Genome Research* 8 (3): 186–94.

Ewing, B., L. Hillier, M. C. Wendl, and P. Green. 1998. “Base-Calling of Automated Sequencer Traces Using Phred. I. Accuracy Assessment.” *Genome Research* 8 (3): 175–85.

Ferris, Patrick, Bradley J. S. C. Olson, Peter L. De Hoff, Stephen Douglass, David Casero, Simon Prochnik, Sa Geng, et al. 2010. “Evolution of an Expanded Sex-Determining Locus in Volvox.” *Science* 328 (5976): 351–54.

Filée, Jonathan. 2014. “Multiple Occurrences of Giant Virus Core Genes Acquired by Eukaryotic Genomes: The Visible Part of the Iceberg?” *Virology* 466-467 (October): 53–59.

Fischer, Matthias G. 2021. “The Virophage Family Lavidaviridae.” *Current Issues in Molecular Biology* 40: 1–24.

Gaïa, Morgan, Lingjie Meng, Eric Pelletier, Patrick Forterre, Chiara Vanni, Antonio Fernandez-Guerra, Olivier Jaillon, et al. 2023. “Mirusviruses Link Herpesviruses to Giant Viruses.” *Nature*, April. https://doi.org/10.1038/s41586-023-05962-4.

Gallot-Lavallée, Lucie, and Guillaume Blanc. 2017. “A Glimpse of Nucleo-Cytoplasmic Large DNA Virus Biodiversity through the Eukaryotic Genomics Window.” *Viruses* 9 (1). https://doi.org/10.3390/v9010017.

Gordon, D., C. Abajian, and P. Green. 1998. “Consed: A Graphical Tool for Sequence Finishing.” *Genome Research* 8 (3): 195–202.

Grigoriev, Igor V., Richard D. Hayes, Sara Calhoun, Bishoy Kamel, Alice Wang, Steven Ahrendt, Sergey Dusheyko, et al. 2021. “PhycoCosm, a Comparative Algal Genomics Resource.” *Nucleic Acids Research* 49 (D1): D1004–11.

Guo, Jiarong, Ben Bolduc, Ahmed A. Zayed, Arvind Varsani, Guillermo Dominguez-Huerta, Tom O. Delmont, Akbar Adjie Pratama, et al. 2021. “VirSorter2: A Multi-Classifier, Expert-Guided Approach to Detect Diverse DNA and RNA Viruses.” *Microbiome* 9 (1): 37.

Hannat, Sihem, Pierre Pontarotti, Philippe Colson, Marie-Line Kuhn, Eric Galiana, Bernard La Scola, Sarah Aherfi, and Franck Panabières. 2021. “Diverse Trajectories Drive the Expression of a Giant Virus in the

Oomycete Plant Pathogen Phytophthora Parasitica.” *Frontiers in Microbiology* 12 (June): 662762.
Huse, Susan M., Julie A. Huber, Hilary G. Morrison, Mitchell L. Sogin, and David Mark Welch. 2007. “Accuracy and Quality of Massively Parallel DNA Pyrosequencing.” *Genome Biology* 8 (7): R143.
Iyer, L. M., L. Aravind, and E. V. Koonin. 2001. “Common Origin of Four Diverse Families of Large Eukaryotic DNA Viruses.” *Journal of Virology* 75 (23): 11720–34.
Juranek, Stefan A., and Katrin Paeschke. 2012. “Cell Cycle Regulation of G-Quadruplex DNA Structures at Telomeres.” *Current Pharmaceutical Design* 18 (14): 1867–72.
Kuo, Alan, Brian Bushnell, and Igor V. Grigoriev. 2014. “Chapter One - Fungal Genomics: Sequencing and Annotation.” In *Advances in Botanical Research*, edited by Francis M. Martin, 70:1–52. Academic Press.
Lagesen, Karin, Peter Hallin, Einar Andreas Rødland, Hans-Henrik Staerfeldt, Torbjørn Rognes, and David W. Ussery. 2007. “RNAmmer: Consistent and Rapid Annotation of Ribosomal RNA Genes.” *Nucleic Acids Research* 35 (9): 3100–3108.
Leonard, Guy, Aurélie Labarre, David S. Milner, Adam Monier, Darren Soanes, Jeremy G. Wideman, Finlay Maguire, et al. 2018. “Comparative Genomic Analysis of the ‘Pseudofungus’ Hyphochytrium Catenoides.” *Open Biology* 8 (1). https://doi.org/10.1098/rsob.170184.
Li, Xianran, Chengsong Zhu, Zhongwei Lin, Yun Wu, Dabao Zhang, Guihua Bai, Weixing Song, et al. 2011. “Chromosome Size in Diploid Eukaryotic Species Centers on the Average Length with a Conserved Boundary.” *Molecular Biology and Evolution* 28 (6): 1901–11.
Messyasz, Adriana, Stephanie M. Rosales, Ryan S. Mueller, Teresa Sawyer, Adrienne M. S. Correa, Andrew R. Thurber, and Rebecca Vega Thurber. 2020. “Coral Bleaching Phenotypes Associated With Differential Abundances of Nucleocytoplasmic Large DNA Viruses.” *Frontiers in Marine Science* 7. https://doi.org/10.3389/fmars.2020.555474.
Moniruzzaman, Mohammad, Carolina A. Martinez-Gutierrez, Alaina R. Weinheimer, and Frank O. Aylward. 2020. “Dynamic Genome Evolution and Complex Virocell Metabolism of Globally-Distributed Giant Viruses.” *Nature Communications* 11 (1): 1710.
Moniruzzaman, Mohammad, Alaina R. Weinheimer, Carolina A. Martinez-Gutierrez, and Frank O. Aylward. 2020. “Widespread Endogenization of Giant Viruses Shapes Genomes of Green Algae.” *Nature* 588 (7836): 141–45.
Murakoshi, Yumi, Takayuki Shimeki, Daiske Honda, and Yoshitake Takao. 2021. “Draft Genome Sequence of Sicyoidochytrium Minutum DNA Virus Strain 001.” *Microbiology Resource Announcements* 10 (23): e0041821.
Paeschke, Katrin, Stefan Juranek, Tomas Simonsson, Anne Hempel, Daniela Rhodes, and Hans Joachim Lipps. 2008. “Telomerase Recruitment by the Telomere End Binding Protein-β Facilitates G-Quadruplex DNA Unfolding in Ciliates.” *Nature Structural & Molecular Biology* 15 (6): 598–604.
Pollak, A. 1979. “Abnormal Development Of Thraustochytrium Aureum, A Biflagellate Marine Phycomycete.” Ph.D., City University of New York. https://search.proquest.com/openview/7bb6f5b20ee6ee4e30bf269d96461572/1?pq-origsite=gscholar&cbl=18750&diss=y&casa_token=V6zxCV4DdDYAAAAA:Hp8Tam9jvPAQ8kHssjxdRemA9xzUZisGqMQ0lxjzVSAHagp0-ZL7r3GBip0xoD6hUt8ddlG-bA.
Rius, Mariana, Joshua S. Rest, Gina V. Filloramo, Anna M. G. Novák Vanclová, John M. Archibald, and Jackie L. Collier. 2023. “Horizontal Gene Transfer and Fusion Spread Carotenogenesis among Diverse Heterotrophic Protists.” *Genome Biology and Evolution*, February. https://doi.org/10.1093/gbe/evad029.
Silva, Genivaldo Gz, Bas E. Dutilh, T. David Matthews, Keri Elkins, Robert Schmieder, Elizabeth A. Dinsdale, and Robert A. Edwards. 2013. “Combining de Novo and Reference-Guided Assembly with Scaffold_builder.” *Source Code for Biology and Medicine* 8 (1): 23.
Takao, Y., K. Nagasaki, and D. Honda. 2007. “Squashed Ball-like dsDNA Virus Infecting a Marine Fungoid Protist Sicyoidochytrium Minutum (Thraustochytriaceae, Labyrinthulomycetes).” *Aquatic Microbial Ecology: International Journal* 49 (November): 101–8.
Takao, Yoshitake, Kazuyuki Mise, Keizo Nagasaki, Tetsuro Okuno, and Daiske Honda. 2006. “Complete Nucleotide Sequence and Genome Organization of a Single-Stranded RNA Virus Infecting the Marine Fungoid Protist Schizochytrium Sp.” *The Journal of General Virology* 87 (Pt 3): 723–33.
Takao, Yoshitake, Keizo Nagasaki, Kazuyuki Mise, Tetsuro Okuno, and Daiske Honda. 2005. “Isolation and Characterization of a Novel Single-Stranded RNA Virus Infectious to a Marine Fungoid Protist,

Schizochytrium Sp. (Thraustochytriaceae, Labyrinthulea).” *Applied and Environmental Microbiology* 71 (8): 4516–22.

Takao, Yoshitake, Yuji Tomaru, Keizo Nagasaki, and Daiske Honda. 2015. “Ecological Dynamics of Two Distinct Viruses Infecting Marine Eukaryotic Decomposer Thraustochytrids (Labyrinthulomycetes, Stramenopiles).” *PloS One* 10 (7): e0133395.

Wang, Feng, Ming-Liang Tang, Zhi-Xiong Zeng, Ren-Yi Wu, Yong Xue, Yu-Hua Hao, Dai-Wen Pang, Yong Zhao, and Zheng Tan. 2012. “Telomere- and Telomerase-Interacting Protein That Unfolds Telomere G-Quadruplex and Promotes Telomere Extension in Mammalian Cells.” *Proceedings of the National Academy of Sciences of the United States of America* 109 (50): 20413–18.
